# Emergence and global spread of *Listeria monocytogenes* main clinical clonal complex

**DOI:** 10.1101/2020.12.18.423387

**Authors:** Alexandra Moura, Noémie Lefrancq, Alexandre Leclercq, Thierry Wirth, Vítor Borges, Brent Gilpin, Timothy J. Dallman, Joachim Frey, Eelco Franz, Eva M. Nielsen, Juno Thomas, Arthur Pightling, Benjamin P. Howden, Cheryl L. Tarr, Peter Gerner-Smidt, Simon Cauchemez, Henrik Salje, Sylvain Brisse, Marc Lecuit, for the *Listeria* CC1 Study Group

## Abstract

Retracing microbial emergence and spread is essential to understanding the evolution and dynamics of pathogens. The bacterial foodborne pathogen *Listeria monocytogenes* clonal complex 1 (*Lm*-CC1) is the most prevalent clonal group associated with listeriosis, and is strongly associated with cattle and dairy products. Here we analysed 2,021 *Lm*-CC1 isolates collected from 40 countries, since the first *Lm* isolation to the present day, to define its evolutionary history and population dynamics. Our results suggest that *Lm*-CC1 spread worldwide from North America following the Industrial Revolution through two waves of expansion, coinciding with the transatlantic livestock trade in the second half of the 19^th^ century and the rapid growth of cattle farming in the 20^th^ century. *Lm*-CC1 then firmly established at a local level, with limited inter-country spread. This study provides an unprecedented insight into *Lm*-CC1 phylogeography and dynamics and can contribute to effective disease surveillance to reduce the burden of listeriosis.

---

*Listeria monocytogenes* (*Lm*) is a foodborne bacterial zoonotic pathogen that can cause listeriosis, a severe infection with a high case-fatality rate in immunocompromised individuals^1,2^. Molecular studies have shown the clonal population structure of *Lm*^3,4^ and the worldwide distribution of clonal complex 1 (*Lm*-CC1, initially called epidemic clone ECI^5,6^), a serotype 4b cosmopolitan clonal group defined by multilocus sequence typing (MLST), which was first isolated from an Italian soldier with meningitis during the first world war (WWI)^7,8^. Interestingly, *Lm*-CC1 has been reported as the most prevalent clinical clonal complex in several countries^9–14^, and data collected on NCBI Sequence Read Archive also support this conclusion (**Supplementary Figure S1**).

While there is no inter-human transmission of listeriosis, it was only in the mid 1980’s that the foodborne origin of human listeriosis was formally proven^15^. Since then, *Lm*-CC1 has been reported in different food matrixes, including dairy products^16–18^ which can be heavily contaminated^19^ and constitute a major source of human listeriosis^20,21^. Previous studies have also demonstrated the hypervirulence of *Lm*-CC1^9^, and its higher efficiency in gut colonization and fecal shedding, compared to hypovirulent *Lm* clones^16,17,22,23^. Moreover, increasing evidence suggests that cattle, which are frequent *Lm* asymptomatic carriers^24–28^ and contribute to *Lm* enrichment in soils^25^, may constitute a reservoir for *Lm*-CC1. In addition to *Lm* subclinical infections that may contaminate milk^23,26^, the long-term persistence of *Lm* in cattle manure-amended soils^29^ also poses serious risks of transmission to fresh produce.

Understanding the global evolution of *Lm*-CC1, which is now spread over all continents^6^, as well as its emergence and dissemination across different spatial levels is critical to understand *Lm* population dynamics and to develop better control strategies, especially in countries with ageing and/or immunosuppressed populations who are most at risk for severe infection. However the complex movement of livestock and food products associated with asymptomatic intestinal colonization complicates traditional epidemiological investigations aimed to decipher *Lm* epidemiology by linking isolates in space and time.

Here we took a population biology approach to fill this knowledge gap and conducted the largest genomic *Lm*-CC1 study to date, combining genomic and evolutionary approaches to decipher its evolutionary history and pattern of emergence and spread.

## Results

### *Lm*-CC1 is composed of 3 sublineages of uneven prevalence

We analyzed 2,021 genomes, including 1,230 newly sequenced isolates, originating from 40 countries in 6 continents and diverse sources (**Figure 1a**; **Supplementary Table S1**). We covered a time span of 98 years, from the first *Lm* isolation to the present time (1921-2018), and included all contemporary clinical isolates collected between 2012 and mid-2017 within the surveillance framework of 7 countries over 3 continents (**Figure 1a,b**).

**Figure 1.**
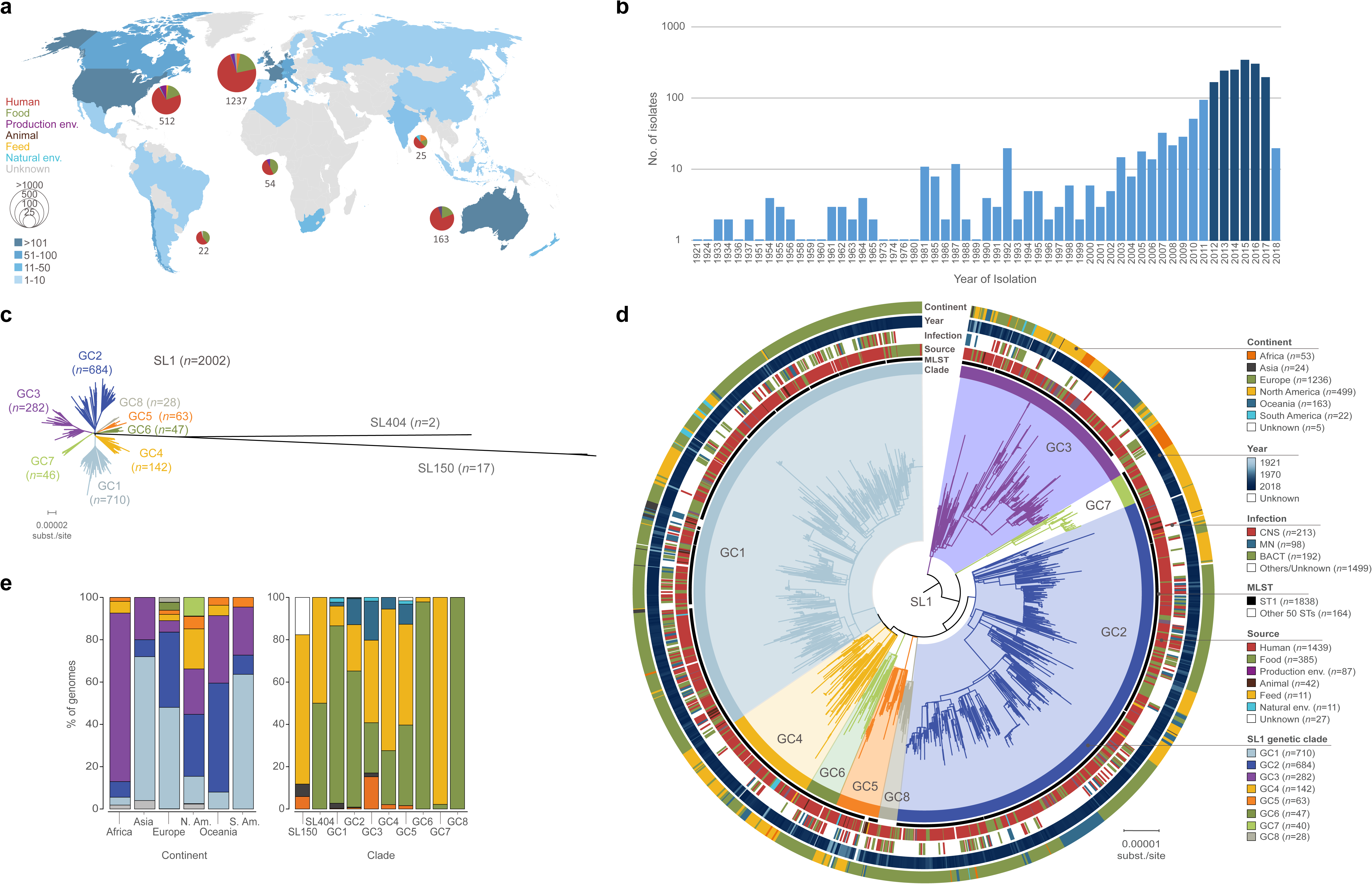
Geographical and temporal distribution of the isolates used in this study (*N*=2,021) and phylogenetic analyses. **a.** Geographical distribution and source distribution. Sampled countries are colored in blue, with hue gradient according to the number of isolates. Pie charts are proportional to the number of isolates sampled in each continent and represent the repartition of sample source types, using the source color key indicated in panel d. Out of 2,021 genomes, 8 isolates had unknown sampling location and are not shown in the map. Temporal distribution of isolates collected in this study. Darker blue bars indicate the period for which exhaustive clinical sampling was obtained for 7 countries spanning 3 continents (2012-2017; US, FR, UK, DK, NL, AU, NZ). **c.** Unrooted maximum-likelihood phylogenetic tree of 2,021 *Lm*-CC1 genomes. The tree was generated from analysis (GTR+F+G4 model, 1000 ultra-fast bootstraps) of a 1.29 Mb recombination-purged core genome alignment. **d.** Midpoint rooted maximum-likelihood phylogenetic tree of 2,002 SL1 genomes based on a recombination-purged core genome alignment of 1.29 Mb. The four external rings indicate the world region, year, type of infection and source type, respectively. The two inner rings indicate ST1 isolates and the 8 SL1 genetic clades identified in this study, respectively. e. Percentage of genomes by world region (left) and phylogroup (right). Partitions are colored by world regions and phylogroup, using the same color code as in panel d.

*Lm*-CC1 genome sizes ranged from 2.77 to 3.25 Mbp, with an average number of 2,879±77 coding sequences and G+C content of 37.7-38.3% (**Supplementary Figure S2**). On the basis of MLST^4^, 58 sequence types (STs) could be distinguished, with ST1 representing 91% (*n*=1838) of isolates. On the basis of core genome MLST (cgMLST)^30^, we identified within *Lm*-CC1 867 cgMLST types, 92% of which were country-specific (**Supplementary Figure S3**). Rarefaction analysis based on cgMLST resampling did not reach an asymptote (**Supplementary Figure S3**), indicating that despite the high number of sequences obtained in this study, a significant amount of *Lm*-CC1 diversity remains undetected.

To better understand the phylogenetic diversity of *Lm*-CC1, we built maximum likelihood phylogenies and identified 3 sublineages (SL1, SL404 and SL150, named based on their smallest ST number). These sublineages have highly uneven frequency (**Figure 1c,d**; **Supplementary Figure S4**), with SL1 (*n*=2002, isolated worldwide) representing 99.1% of the isolates, while 0.1% are SL404 (*n*=2, found in Europe and North America) and 0.8% represent SL150 (*n*=17, found in North America, Africa and Asia). Within SL1, we further identified 8 distinct genetic clades, which we named GC1 to GC8 by decreasing prevalence (**Figure 1**; **Supplementary Figure S4**). The average genetic distance was 1166±134 wgSNPs (and 478±20 cgMLST alleles) between *Lm*-CC1 sublineages, and 76±16 wgSNPs (and 40±9 cgMLST alleles) within SL1 clades (**Supplementary Table S2**; **Supplementary Figure S5**). The finding that SL1 is by far the major sublineage in *Lm*-CC1 is consistent with either its increased virulence and/or transmission or that SL404 and SL150 are restricted to some yet unknown ecological niches. Within SL1, all different genetic clades were well represented, with strong spatial structure: GC1 is the most prevalent clade in Europe (48%, 593/1237), Asia (68%, 17/25) and South America (64%, 14/22); GC2 is the most prevalent clade in North America (29%, 150/512) and Oceania (52%, 84/163), while GC3 is the most prevalent clade in Africa (80%, 43/54) (**Figure 1e**; **Supplementary Figure S6**).

### The *Lm*-CC1 pangenome is diverse

Analysis of *Lm*-CC1 pangenome identified 10,789 orthologous coding sequences (BlastP identity cut-off of ≥95%), 2,649 of which (92% of the average isolate genome content) present in at least 95% of isolates (core genome) (**Supplementary Figure S7**). The accessory genome included 8,140 gene families, of which 2,844 (35%) were unique to one isolate, and was enriched in transcription, replication/repair and cell wall functions, as well as in gene families of unknown function (**Supplementary Figure S7**). Plasmids were present in 6% (120/2021) of isolates, and were more prevalent in GC7 (83%, **Supplementary Figure S7**). Intact prophages were present in 62% isolates (1263/2021), and were distributed across the breadth of CC1 phylogeny, except in SL404 (**Supplementary Figure S7**). In contrast to *Listeria* pathogenic islands LIPI-1^31^ and LIPI-3^32^ which were present in all isolates, the *Listeria* genomic island LGI2-1^33^, previously identified in CC1 isolates encoding resistance to cadmium and arsenic, was present in 14% (277/2021) isolates and only in GC3 (80%, 225/283), GC5 (60%, 38/63) and SL150 (82%, 14/17; **Supplementary Figure S7**). Sublineage-specific genes were detected (*n*=81; **Supplementary Tables S3 and S4**) and pangenome-wide association analyses identified 24 genes that are associated with a clinical origin (**Supplementary Table S5**). The impact of these traits on isolates’ differential ecology or virulence remains to be studied, yet the presence of human isolates in all sublineages and clades shows that pathogenic isolates are not restricted to a specific *Lm*-CC1 clade.

### Emergence and worldwide spread of *Lm*-CC1 main sublineage (SL1) occurred in the last 200 years

To understand *Lm*-CC1 evolution and spread, we performed temporal and phylogeographic analyses on a subset of 200 genomes representative of *Lm*-CC1 genetic and geographic diversity using BEAST^34^, and on the full dated dataset (1,972 *Lm*-CC1 genomes) using Treedater^35^ (**Supplementary Figures S8 and S9**) and PastML^36^, under an uncorrelated relaxed clock model (see Material and Methods for details). We estimate a core genome substitution rate of 1.95×10^−7^ substitutions/site/year (95% CI: 1.75×10^−7^−2.15×10^−7^; **Supplementary Figure S8**), consistent with previous findings^30^. We estimate that *Lm*-CC1 originated about 1,800 years ago (date: 197 AD; 95% CI: 860 BC – 1045 AD; **Figure 2b**) and infer that its last common ancestor evolved in North America (**Supplementary Figure S10**), long before European colonization and the introduction of cattle in the Americas at the end of the 15^th^ century^37^. Even though the low number of genomes available for Asia, Africa and South America could bias this estimation, the estimated origin was also supported by the measures of population variability, which showed higher genetic diversity within North America (**Supplementary Figure S5**; **Supplementary Table S2**), and by the basal position of North American *Lm*-CC1 isolates in the phylogeny (**Figure 2b, Supplementary Figure S10**). Whether *Bison bison* populations, which are phylogenetically and ecologically related to bovine and dominated North American prairies prior to colonization by the Europeans and their livestock, played a role in its dispersion remains unknown.

**Figure 2.**
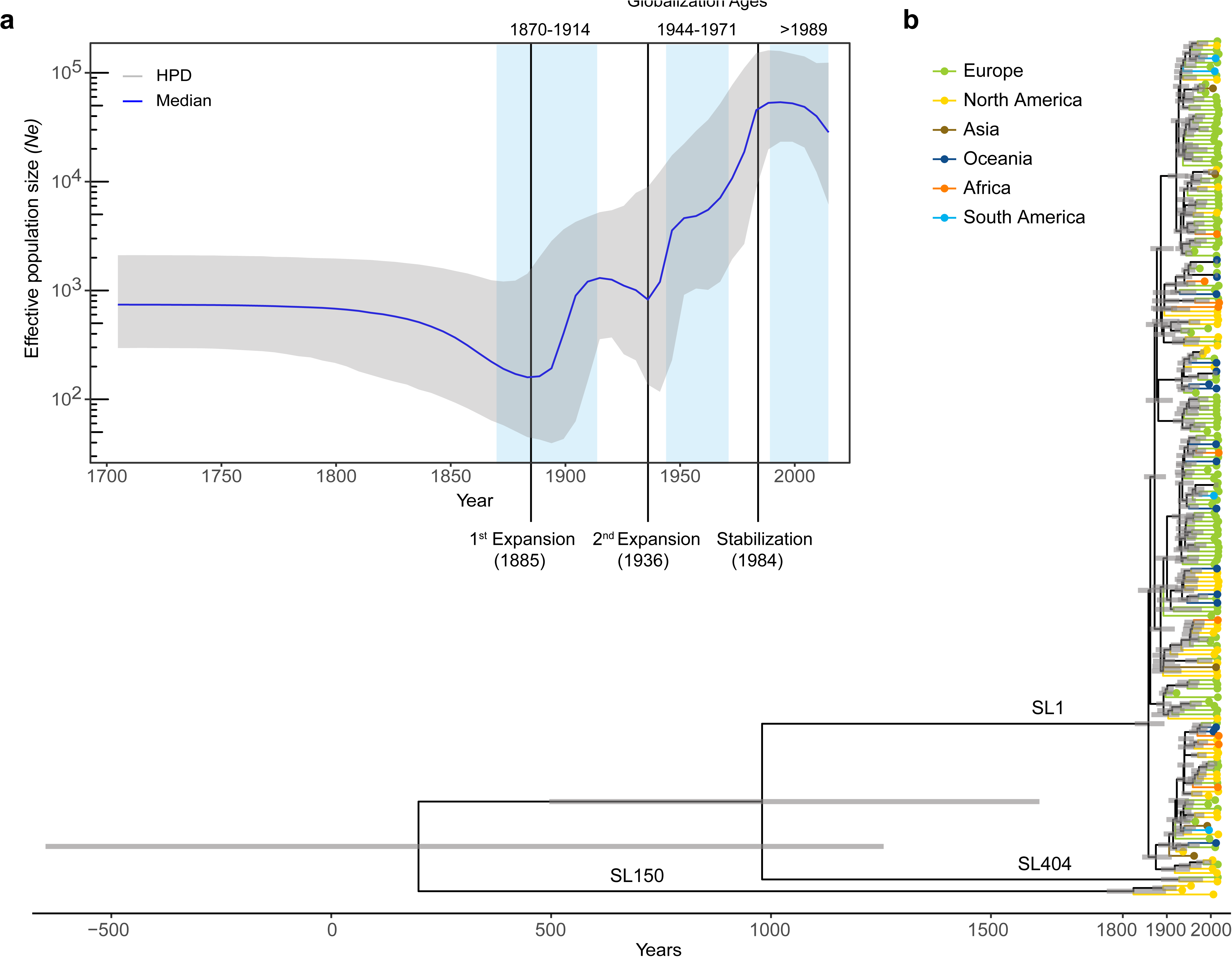
Bayesian temporal and demographic analyses on a representative 200 isolate dataset. **a.** Bayesian skyline plot (BSP) with the estimation of *Lm*-CC1 effective population size (*Ne*). The y-axis refers to the predicted number of individuals (log scale) and the x-axis to the timescale (in years). The median population size is marked in blue with its 95% high posterior density (HDP) in gray. Blue vertical panels delimitate the three globalization ages (1870-1914, 1944-1971, 1989-present). **b.** Bayesian time-calibrated tree. Nodes represent the estimated mean divergence times and gray bars represent the 95% HPD confidence intervals of node age. Scale indicates time (in years). Terminal branches and tips are colored by continents, as indicated in the key panel.

Demographic analyses performed using the Bayesian Skyline Plot method^38^ (**Figure 2a**) show that *Lm*-CC1 effective population size was stable up to the middle of the 19^th^ century, followed by two waves of expansion: the first in the late 1880s and the second in the 1930s, coinciding with the first and second ages of globalization, respectively. Tajima’s D statistic^39^ also supported a recent CC1 population expansion and SL1 emergence (D<0; **Supplementary Table S2**). SL1 emerged in North America approximately 160 years ago (date: 1859, 95% CI: 1821-1889), thus closely following the start of the Industrial Revolution (**Figure 3**). The first SL1 introductions into Europe occurred around 1868 (GC6/GC8 ancestor, 95% CI: 1827-1890), 1871 (GC3/GC7 ancestor, 95% CI: 1838-1905) and 1889 (GC2, 95% CI: 1852-1909), concomitant with the 1870 North Atlantic Meat trade agreement^40^. Under this agreement, surplus cattle in North America were shipped to Europe, which had experienced severe livestock shortages due to widespread disease outbreaks (contagious bovine pleuropneumonia and foot and mouth disease), leading to an unprecedented man-made 1000-fold increase in cattle movement From North America to Europe^41^. Within the same period, intra-continental diversification also took place, likely driven by cattle movements across North America and railway expansion in North America and Europe. The first SL1 introductions that occurred in Oceania (1903, GC2) followed the ‘Great Drought’ of 1895-1903, which severely affected livestock^42^.

**Figure 3.**
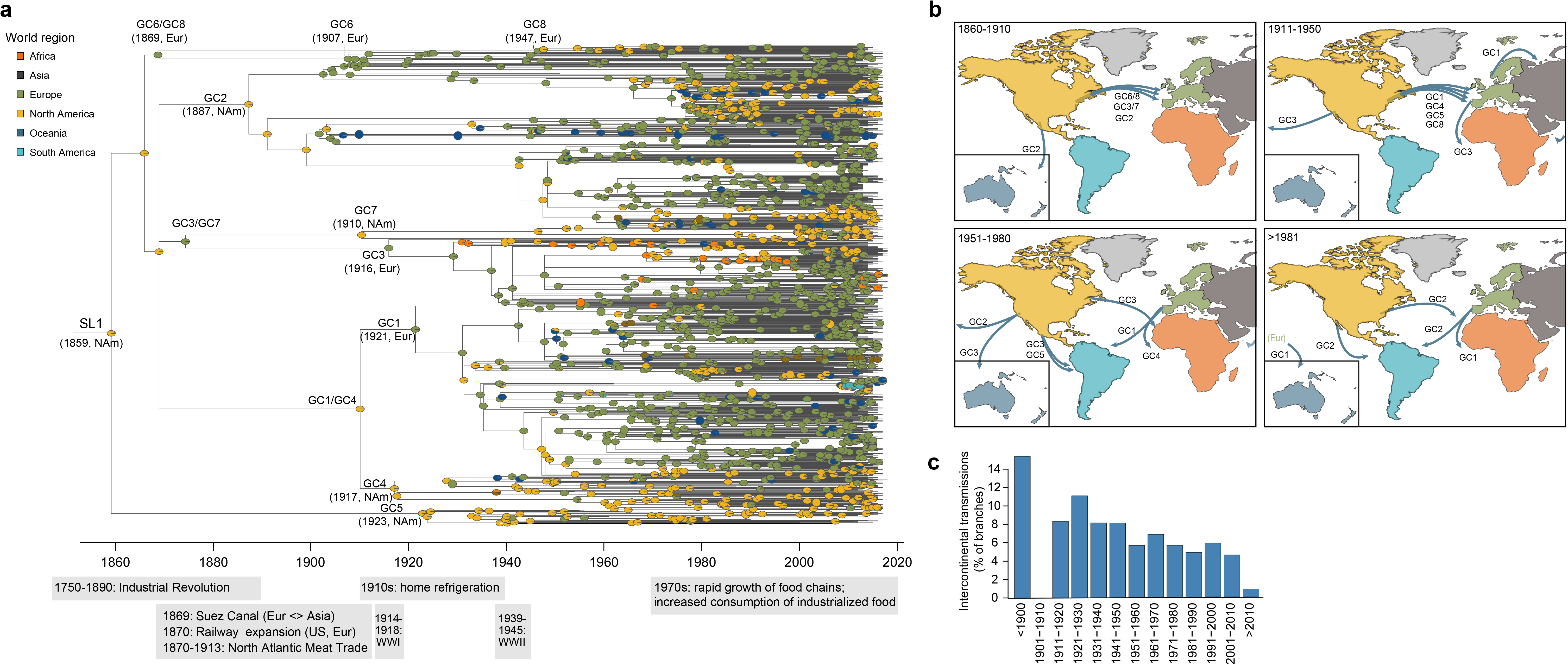
Phylogeography of sublineage SL1. **a.** Time-calibrated phylogeny based on the 1956 SL1 genomes. Pies at the nodes represent the probability of ancestral geographical locations, estimate using PastML using the MPPA method with an F81-like model. **b.** Inferred spread of SL1 populations across continents. The first introductions of each phylogroup are represented by arrows from their estimated world region origin. **c.** Proportion of inter-continental transitions per 10-year bins, normalized by the total number of phylogenetic branches per bin.

In the following decades and after WWI, multiple CC1 introductions continued from North America into Europe (GC1, GC4, GC5 and GC8) and Asia (GC3) and from Europe to Africa (GC3) (**Figure 3a-b**). The rate of intercontinental bacterial movement declined after 1930s (**Figure 3c**), concomitant with the protectionist trade policies that followed the ‘Great Depression’, which led to a sharp reduction of livestock exports from the USA during the first half of the 20^th^ century^43^. A second wave of SL1 expansion occurred after this period, likely driven by a new increase in intercontinental movements favoured by the industrialization of food production and globalization of the food and cattle trades (**Figures 2a**; **Supplementary Figure S11**). Other important human pathogens that have a zoonotic reservoir such as *Escherichia coli* O157:H7^44^ and *Campylobacter jejuni* ST61^45^, have been estimated to have most recent common ancestors (MRCA) at similar times and to have undergone population expansions in the context of animal trade or intensive cattle farming, respectively.

A stabilization and relative decline of *Lm*-CC1 population is observed after 1984 **(Figure 2a)**, coincident with the major advances in infectious diseases’ prevention in dairy cattle^46^ and with the relative decrease of the dairy cattle population in Western countries, in particular Europe (**Supplementary Figure S11**). It also coincides with the time when human listeriosis foodborne origin was formally proven^15^, which led to the implementation of surveillance programs in North America and Europe^47–50^, in particular in the dairy sector following cheese and milk related *Lm*-CC1 outbreaks^51^. Whether these findings can be observed in other dairy-associated *L. monocytogenes* clonal complexes, such as CC6 (lineage I) or CC37 and CC101 (lineage II)^17,52^ will deserve future studies.

### Recent SL1 transmission chains are mostly local

To further analyze more recent strain transmission dynamics, we compared the genetic diversity of SL1 isolates from 2010-2018 (*n*=1,266) across different spatial scales. To avoid oversampling isolates from outbreak investigations, we excluded all non-clinical isolates from confirmed outbreaks (*n*=91 isolates from 19 outbreaks). We find that pairs of isolates present within the same 2-year period and the same country are 18.7 times (95% CI: 4.7-190.7) more likely to have their MRCA within the past 5 years than pairs of isolates coming from other intra-continental countries >1,000 km apart (**Figure 4a**). Furthermore, we observe no difference in the probability of having a recent MRCA in isolates coming from nearby intracontinental countries (<1,000km) than from further apart. Isolates coming from different continents are about 100 times less likely to have an MRCA within the past 5 years (0.2; 95% CI: 0.01-2.9) than isolates from the same countries (18.7; 95% CI: 4.7-190.7) (**Figure 4a**). This strong local spatial structure persists for very long time periods, with complete mixing of isolates within a continent appearing only after 50 years (**Figure 4a**). At a finer spatial scale, available for France (“*départements*”, sub-regional administrative division in France, **Supplementary Figure S12**), a strong local spatial structure is also evident, with the proportion of genetically close pairs of clinical cases being higher between isolates coming from the same French department (4.4%, 95% CI: 1%-10.6%) than between isolates coming from different departments (0.2%, 95% CI 0.04%-0.5%), with no effect of distance between them (**Figure 4b**). As expected, in densely urban areas with no farming, such as the city of Paris, clinical strains are significantly less likely to share a recent MRCA than in rural areas or other departments (0.0%, 95% CI: 0.0%-4.4% *vs*. 3.9%, 95% CI: 1.0%-9.5%) (**Figure 4c**). This result is consistent with urban infections being driven by unrelated *Lm* introductions originating from across the country. Spatial dependence between French isolates persists for 20 years (**Supplementary Figure S13**), with on average 20 (1/0.05) different sources of human infection present at any one time per department (**Figure 4b**).

**Figure 4.**
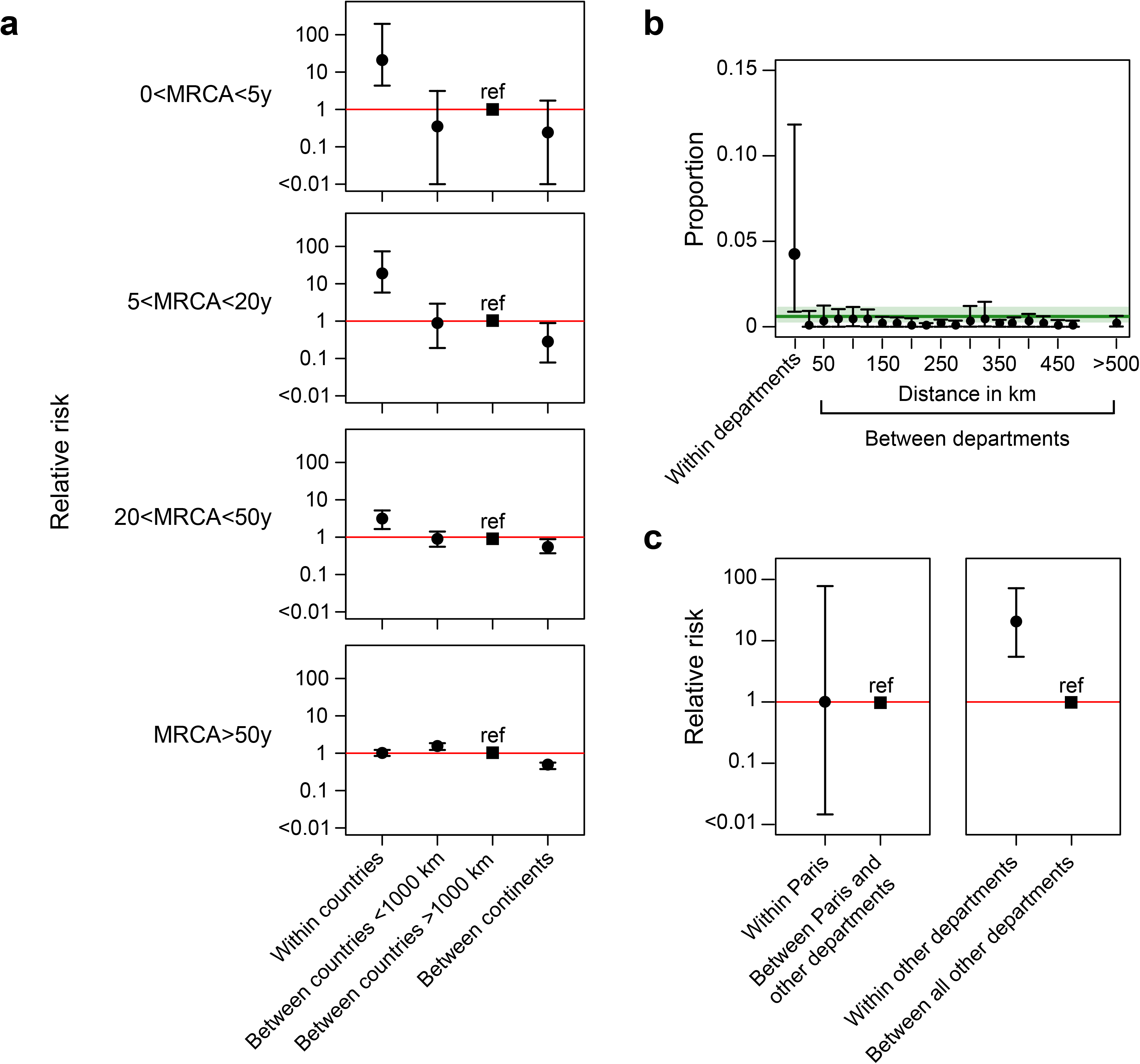
Transmission dynamics of sublineage SL1. **a.** Each point summarizes the relative risk that a pair of isolates has a MRCA within a defined timeframe and between different spatial scales (within the same country, within the same continent or within different continents), relative to the risk that a pair of isolates from countries separated by >1000km have a MRCA in the same range (set as the reference value, ‘ref’). Error bars represent the 95% confidence intervals, based on 100 bootstrap time-calibrated trees. **b.** Proportion of pairs of isolates within the same country (France) sharing a MRCA of 5 or less years in function of the spatial distance within and between administrative departments (shown in the map). The green line indicates the mean proportion of genetically close strains regardless the geographical location. **c.** Left: relative risk for a pair of isolates to share a MRCA of 5 or less years when both are coming from Paris to when coming from another department (*p*=0.43). Right: relative risk for a pair of isolates to share a MRCA of 5 or less years when coming from the same department in France, except Paris, compared to when coming from different departments (*p*<0.001, see Material and Methods for details).

## Discussion

Understanding pathogen evolutionary history is essential to understand the population dynamics and biodiversity of microbial infectious agents, and for effective disease surveillance. Here, we have shown that *Lm*-CC1 has spread worldwide following the Industrial Revolution, and that genotypes are now firmly established at a local level, with decades-long localized persistence. These results are consistent with the establishment of separate, locally entrenched sources of *Lm*-CC1 with limited flow of bacteria either within or between countries, in line with cgMLST analyses in which 92% of clusters are country-specific.

In the absence of inter-human transmission, this observation likely represents persistent infection sources, *i.e*. individual herds and/or production facilities, in which *Lm* can reside for several years^28,53^. Outbreak investigations performed at local scale, including in farm environments, would therefore likely improve the identification of contaminating sources, which remain unknown in about 80% of clusters of human cases^54^. Identifying and eradicating sources along the food chain, from the farm to the fork, could lead to significant long-term reductions in the transmission of the *Lm*-CC1.

The current scarcity of genomes available for Asia, Africa and South America, and from natural and animal reservoirs may overlook other CC1 clades and could have biased our phylogeographic analyses. Nevertheless, this study sheds unprecedented light onto the evolutionary history, epidemiology and population dynamics of *Lm*-CC1. Similar approaches targeting other major globally distributed clonal complexes will allow clarifying their transmission dynamics and uncovering epidemiological specificities of *Lm* clones. Deciphering the dynamics and drivers of *Lm* sublineages across time and space will inform infection control policies and ultimately reduce the burden of listeriosis.

## Methods

### Bacterial isolates and genome sequencing

A total of 2,021 high quality *Listeria monocytogenes* clonal complex 1 (CC1) genomes collected by this study group (*n=*1,230) and from NCBI repositories (*n=*791, as of 14 March 2018) were analyzed. These were part of an initial dataset of 2,154 CC1 genomes, from which 133 were discarded due to low sequencing coverage (<40X after read trimming, *n=*62) or low assembly quality (>200 contigs and/or N50<20Kb, *n=*71)^30^. The 2,021 isolates originated from human (*n*=1,453; 72%) and animal hosts (*n=*44; 2%), food (*n=*387; 19%), food-processing environments (*n=*88; 4%), feed (*n=*11; 0.5%), natural environments (*n=*11; 0.6%) or from unknown sources (*n=*27; 1%) (**Figure 1; Table S1**). Isolates were sampled in 40 countries from 6 continents, between 1921 and 2018 (**Figure 1; Table S1**). Between 2012 and mid-2017, exhaustive sampling was obtained for 7 countries in 3 continents in the context of listeriosis national surveillance programs in Australia (*n=*75), Denmark (*n=*42), France (*n=* 395), The Netherlands (*n=*53), New Zealand (*n=*34), the United Kingdom (*n=*106) and the United States (*n=* 317). Sequencing reads were obtained using Illumina sequencing platforms (Illumina, San Diego, US) and 2×50 bp (*n=*110), 2×75 bp (*n=*2), 2×100 bp (*n=*233), 2×125 bp (*n=*9), 2×150 bp (*n=*1,145), 2×250 bp (*n=*351), 2×300 bp (*n=*138) paired-end runs (**Table S1**).

### Sequence analysis

Whole genome sequencing reads were available for 1,988 out of 2,021 isolates. Reads were trimmed from adapter sequences and non-confident bases using AlienTrimmer v.0.4^55^ (minimum read length of 30 bases and minimum quality Phred score 20, i.e. 99% base call accuracy) and corrected with Musket v.1.1^56^, implemented in fqCleaner v.3.0 (Alexis Criscuolo, Institut Pasteur). FastQC v.0.11.5^57^ was used to assess sequence quality before and after trimming. Assemblies were obtained from paired-ended trimmed reads ≥75 bp (*n=*1,878 isolates) by using SPAdes v.3.11.0^58^ with the automatic k-*mer*, --only-assembler and --careful options. For paired-ended trimmed reads of 50 bp (*n=*111), assemblies were built using CLC Assembly Cell v.5.0.0 (Qiagen, Denmark), with estimated library insert sizes ranging from 50 to 850 bp. Contigs smaller than 500 bp were discarded from both SPAdes and CLC generated assemblies.

### Pangenome analysis

Gene prediction and annotation was carried out from the draft assemblies using Prokka v.1.12^59^. Functional classification was carried out with EGGnog-mapper v2^60^ using DIAMOND (Double Index Alignment of Next-generation sequencing Data)^61^. The presence of plasmids, intact prophages and *Listeria* genomic regions was inferred from the assemblies using MOB-suite v.2.0.1^62^, PHASTER (https://phaster.ca/)^63^ and BIGSdb-*Lm* (http://bigsdb.pasteur.fr/listeria/)^30,64^, respectively. Pangenome analyses were carried out using Roary v.3.12^65^ with an amino acid identity cut-off of 95% and splitting homologous groups containing paralogs into groups of true orthologs. Venn diagrams were obtained using Venny 2.1 (Oliveros, 2007). Pangenome-wide association analyses were performed using treeWAS v.1.0^66^, to control for phylogenetic structure, using a significance threshold of *p*<10^−5^.

### *In silico* molecular typing

PCR-serogrouping (5 loci)^67^, MLST (7 loci)^4^ and cgMLST (1748 loci)^30^ profiles were extracted from draft assemblies using the BIGSdb-*Lm* platform (http://bigsdb.pasteur.fr/listeria/) as previously described^30^. Profiles were compared using the single linkage clustering method implemented in BioNumerics v.7.6 (Applied-Maths). cgMLST profiles were classified into cgMLST types (CT) and sublineages (SL) using previous defined cut-offs (7 and 150 allelic mismatches, respectively, out of 1748 loci)^30^. Rarefaction curves were computed with vegan v. 2.5-6^68^ R package, estimated with the rarefaction function (Joshua Jacobs, joshuajacobs.org/R/rarefaction) using 100 random samples per point.

### Phylogenetic analyses

Core genome multiple sequence alignments were built from the 1748 cgMLST loci concatenated sequences^30^. Briefly, individual allele sequences were translated into amino acids, aligned separately with MUSCLE v.3.8.31^69^ and back-translated into nucleotide sequence alignment. Concatenation of the 1748 loci alignments resulted in a multiple sequence alignment of 1.57 Mb.

In parallel, whole genome SNP (wgSNP)-based alignments were built from trimmed reads and NCBI assemblies using the Snippy v.4.1.0 pipeline (https://github.com/tseemann/snippy). The closed CC1 genome F2365 (accession no. NC_002973.6), from the 1985 Canadian cheese outbreak^70^ was used as reference in read mapping, resulting in an alignment of 2.29 Mb.

Gubbins v.2.2.0^71^ was used to detect recombination regions in both core and whole-genome alignments, using default parameters and a minimum of 3 base substitutions required to identify recombination. Alignment regions positive for recombination were then completely removed from the original alignments, resulting in recombination-free core- and whole-genome alignments of 1.29 Mb and 2.28 Mb, respectively. Maximum likelihood phylogenies were obtained from the recombination-purged alignments using IQ-tree v.1.6.7.2^72^ under the determined best-fit nucleotide substitution model (GTR+F+G4^73^, as determined by ModelFinder^74^) and ultrafast bootstrapping of 1000 replicates^75^. Trees were visualized and annotated with ggtree v.1.14.6^76^ and iTol v.4.2^77^.

To measure the degree of genetic variation within sublineages, genetic clades and geographic locations, the pairwise allelic and SNP distance matrices were calculated from the cgMLST profiles and multiple sequence alignments, respectively. SNP distances were computed taking into account only the ATGC polymorphic positions, extracted from the alignments using SNP-sites v.2.4.1^78^.

The nucleotide diversity and the Tajima’s D statistics per alignment were calculated using the R package PopGenome v.2.6.1^79^.

### Demographic and spatio-temporal analysis

To infer the population size changes, Bayesian skyline plots were obtained with BEAST v1.10.4^34^. The coalescent Bayesian skyline model was chosen due to its flexibility to allow a wide range of demographic scenarios, avoiding the biases of pre-specified parametric models in the estimates of demographic history^38^. Analyses were performed on a random subset of 200 isolates selected out a subset of 422 isolates representative of genomic and geographic diversity of the full dataset (1 isolate per country per cluster of 99% core genome similarity). Sampling times were positively correlated with the genetic divergence (*p*<0.05, F-Statistic test; Supplementary Figure S6), as observed using TempEst v1.5.1^80^. BEAST estimations were made using the nucleotide evolutionary model GTR+Γ4 and a default gamma prior distribution of 1, under an uncorrelated relaxed clock model, to allow each branch of the phylogenetic tree to have its own evolutionary rate^81^. Runs were performed in triplicates, each consisting of MCMC chains of 400 million iterations, with a 25% burn-in. Parameter values were sampled every 10,000 generations. The effective sample size (ESS) values were confirmed to be higher than 200 for all parameters using Tracer v.1.7^82^. The time of the most recent common ancestor (MRCA) and 95% highest posterior densities (95% HPDs) were inferred from the nodes of the maximum clade credibility tree. To assess the significance of the temporal signatures observed, 10 randomized tip date datasets run under the same parameters were used as controls^83^. To assess the robustness of the population size inference to changes in the dataset, a second non-overlapping subset of 200 genomes obtained from the same representative subset of 422 isolates was analyzed using BEAST with the same parameters as described above. Estimations of the effective population size along the years were computed using Tracer v.1.7^82^.

Phylogeography analyses were then extended to the 1972 CC1 genomes for which country and year of isolation were available. Time-calibrated phylogenies were inferred from the maximum likelihood core genome trees (obtained with IQ-tree, as described above) using either Bactdating v1.0.1^84^, Treetime v0.5.2^85^ or Treedater v0.3.0^35^, assuming a relaxed clock model and the estimated substitution rate of 1.954×10^−7^±2.0152×10^−8^ substitutions/site/year (obtained with BEAST as described above). Cophenetic correlations between BEAST and the three alternative large-scale dating methods were evaluated and better *R^2^* coefficient scores were obtained for Treedater (Supplementary Figure S7). For this reason, the latter dated tree was used in further downstream analyses. Ancestral geographic reconstruction was performed with PastML^36^ using the MPPA method with an F81-like model and estimated ancestral state probabilities were mapped onto the full time-calibrated phylogeny using the R package ape v5.3^86^.

### SL1 global transmission dynamics

To infer the transmission dynamics at a recent time scale (Figure 4a and supplementary Figure S12), we focused on the CC1 main sublineage, and we analyzed the genetic similarity of SL1 isolates from 2010-2018 (*n*=1,266) across different temporal and spatial scales, as described before^87^. To avoid oversampling isolates from outbreak investigations, we excluded all non-clinical isolates from confirmed outbreaks (*n*=91 isolates from 19 outbreaks). We computed the probability **P_l_** that a pair of isolates that satisfy a given location criteria that were sampled within two years of each other had a MRCA in a specific range (0-5 years, 5-20 years, 20-50 years, >50 years), relative to the probability **P_ref_** that a pair isolates), sampled within two years of each other, had an MRCA within that particular range. The location criteria used were: i) within countries (both isolates come from the same country); ii) between countries ≤1000 km (isolates come from distinct countries, separated by less than 1000 km, from the same continent); iii) between countries >1000km (isolates come from distinct countries, separated by more than 1000 km, from the same continent; used as reference); and iv) between continents (isolates come from distinct continents). Spatial relationships between isolates were calculated using the centroid coordinates of the countries or regions of origin.

We estimated these probabilities using:

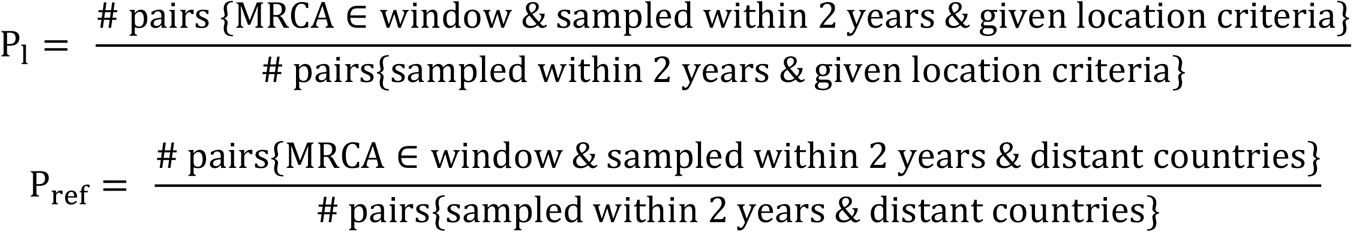

Finally, the relative risk (RR) was given by:

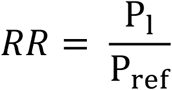

To measure uncertainty, we used a combination of bootstrapping observations and sampling trees from the Treedater v0.3.0 package^35^ to incorporate both sampling and tree uncertainty. Over repeated resamples, we first selected a random tree and calculate the evolutionary distance separating all pairs of sequences. Then, we resampled all the isolates with replacement and recalculate RR each time. The 95% confidence intervals are the 2.5% and 97.5% quantiles from the resultant distribution from 1000 resampling events.

### SL1 local transmission dynamics

To assess the SL1 local transmission dynamics, we used available data from France. We computed the proportion of closely related pairs of French isolates (defined as having a MRCA<5years) as a function of the spatial distance within and between administrative Departments (**Figure 4b**):

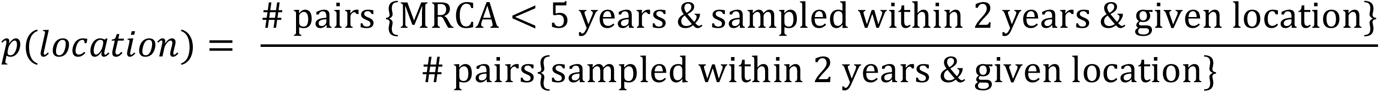

The different location criteria used are: i) within Department: both isolates come from the same Department; ii) between Departments: isolates come from different Departments, separated by a distance from 50 to >500km. The French Departments are shown in the map in **Figure S11.**

As shown in Salje et al.^87^, the reciprocal of *p(within department)* represents the lower limit of the number of sources of human infection circulating within a Department.

To assess uncertainty, we used the bootstrapping approach as described above.

To explore possible differences between Departments, we computed the relative risk that a pair of isolates share a MRCA of less than 5 years when both come from the same department compared to when coming from different departments. We looked at 2 different groups of departments: i) Paris alone **(Figure 4c, left): w**ithin Paris (both isolates come from Paris) and between Paris and other departments (for each pair of isolates, one of them come from Paris, and the other one from another department); ii) other departments, except Paris **(Figure 4c, right)**: with other departments (both isolates come from the same department, excluding Paris) and between all other departments (isolates come from 2 different departments, excluding Paris). For each group, to compute the relative risk *RR*, we used the same approach as explained above. We estimated:

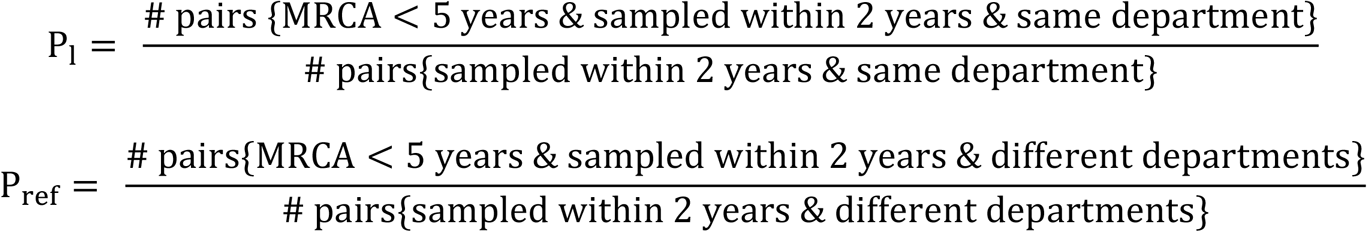

Finally, the relative risk is given by:

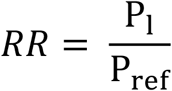

To determine uncertainty, we used the same bootstrapping approach as described above. To assess the statistical significance of each *RR*, we performed a one-tailed test. We set the null hypothesis (*H_0_*) as *RR* ≤ 1, and alternative hypothesis (*H*_1_) as *RR* > 1. For each group, composed *N* bootstrap events, we computed:

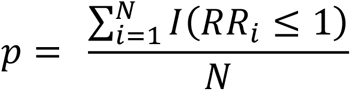

## Data availability

All sequence data will be made available in NCBI-SRA and EBI-ENA public archives upon acceptation.

## Acknowledgements

The findings and conclusions in this report are those of the authors and do not necessarily represent the official position of the Centers for Disease Control and Prevention. The authors thank all participating laboratories and the PulseNet International Network members for their contributions. The authors are also grateful to Martin Wiedmann, Mark Achtman and Jana Haase for providing cultures of historical isolates, to Thomas Cantinelli and Laure Diancourt for contributions to the initial sequencing of CC1 isolates, to Keith Jolley, Youssef Ghorbal and Bryan Brancotte for BIGSdb-*Listeria* maintenance and software updates, to Eduardo Rocha, Etienne Simon-Lorière, Anna Zhukova, Sophie Creno, Eric Deveaud, Guy Bayle and Erik Volz for insightful feedback on methodological issues, and François-Xavier Weill for critical reading. This work used the computational and storage services (TARS cluster) provided by the IT department at Institut Pasteur, Paris.

## Author contributions

ML coordinated the project. ML and SB conceived and designed the study. AM, NL, TW, SC, HS analysed the data, together with SB and ML. AL, VB, BG, TJD, JF, EF, EMN, JT, AP, BPH, CT, PGS, SB, ML and the *Listeria* CC1 study group obtained the isolates, acquired metadata data collection and genome sequences. AM, HS and ML wrote the manuscript. All authors commented and edited the final version of the manuscript.

## SUPPLEMENTARY MATERIAL

### 1. Supplementary tables

**Table S1. Isolates included in this study. [xls]**

**Table S2.**
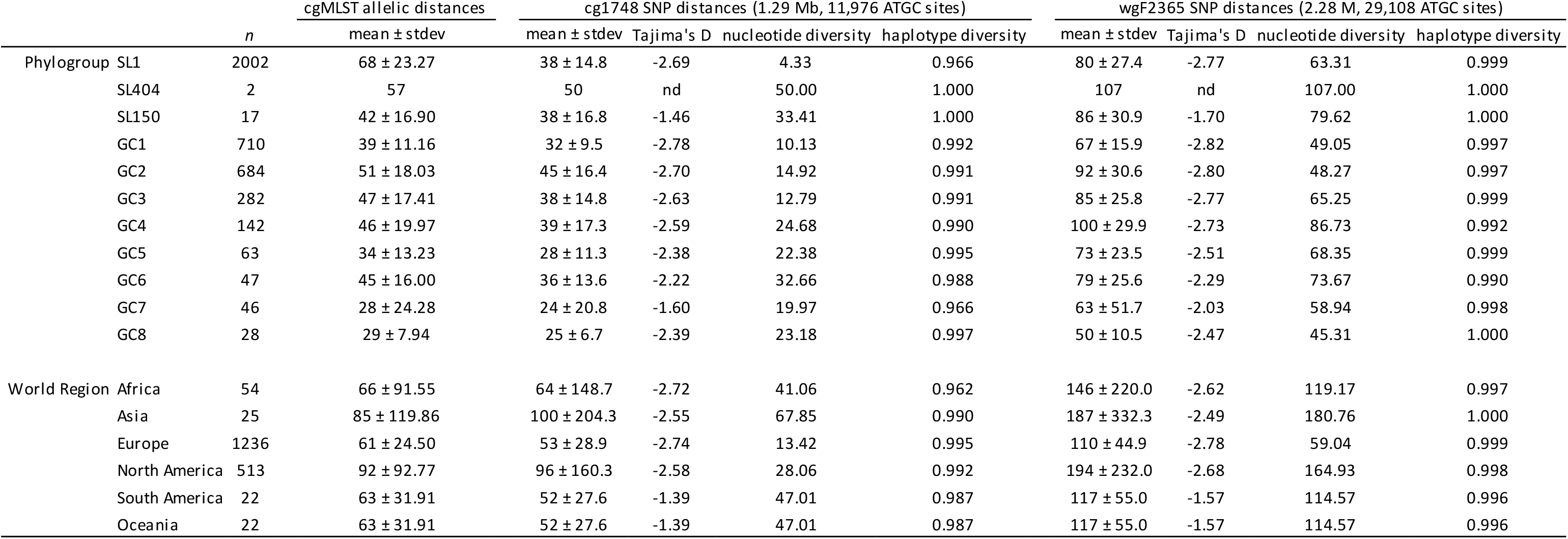
Genetic characteristics of CC1 sublineages and SL1 genetic clades and statistics by world region.

**Table S3.**
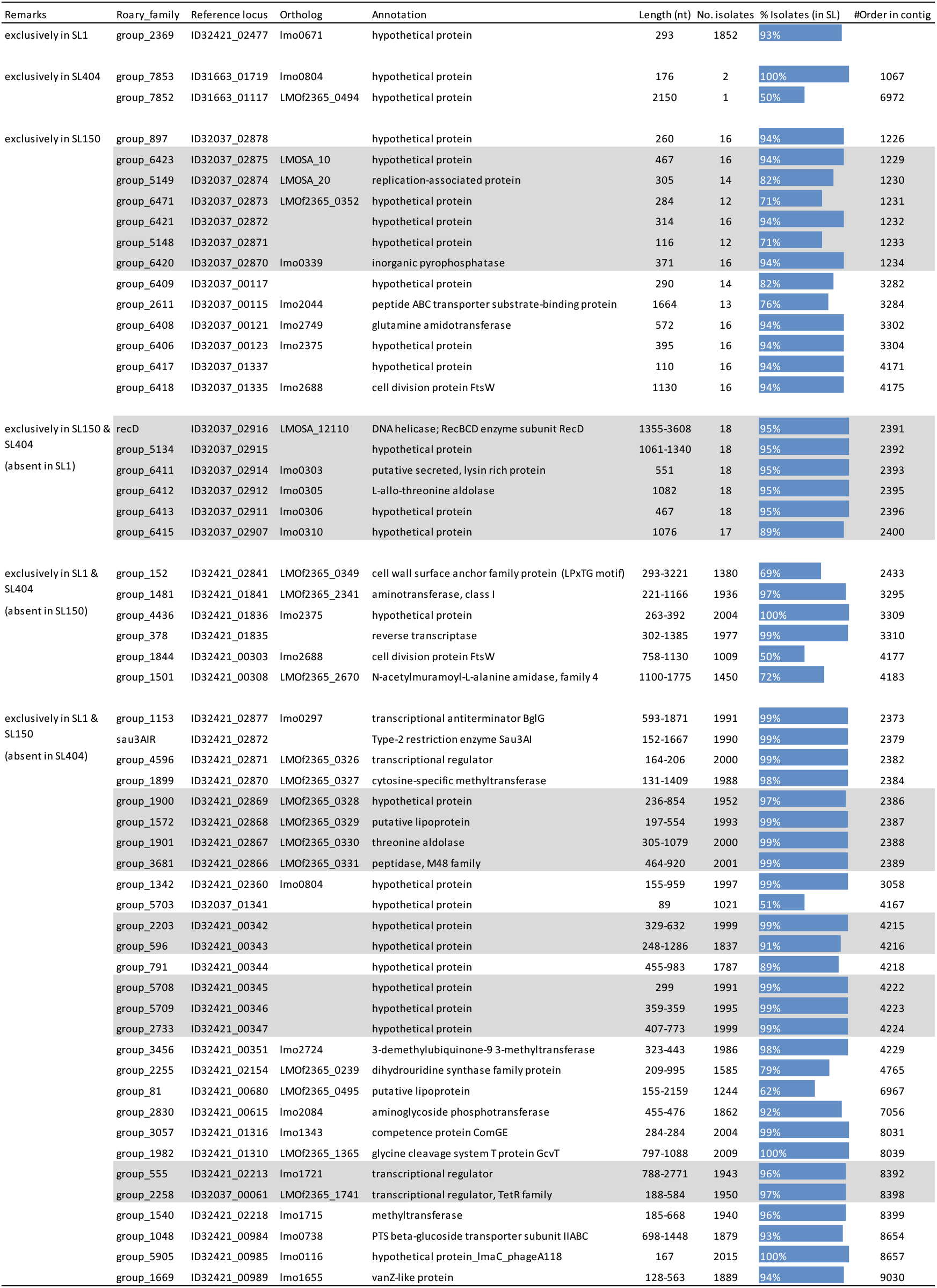
Sublineage-specific genes present in at least 50% of isolates. Gray shades highlight genes within the same genomic context.

**Table S4.**
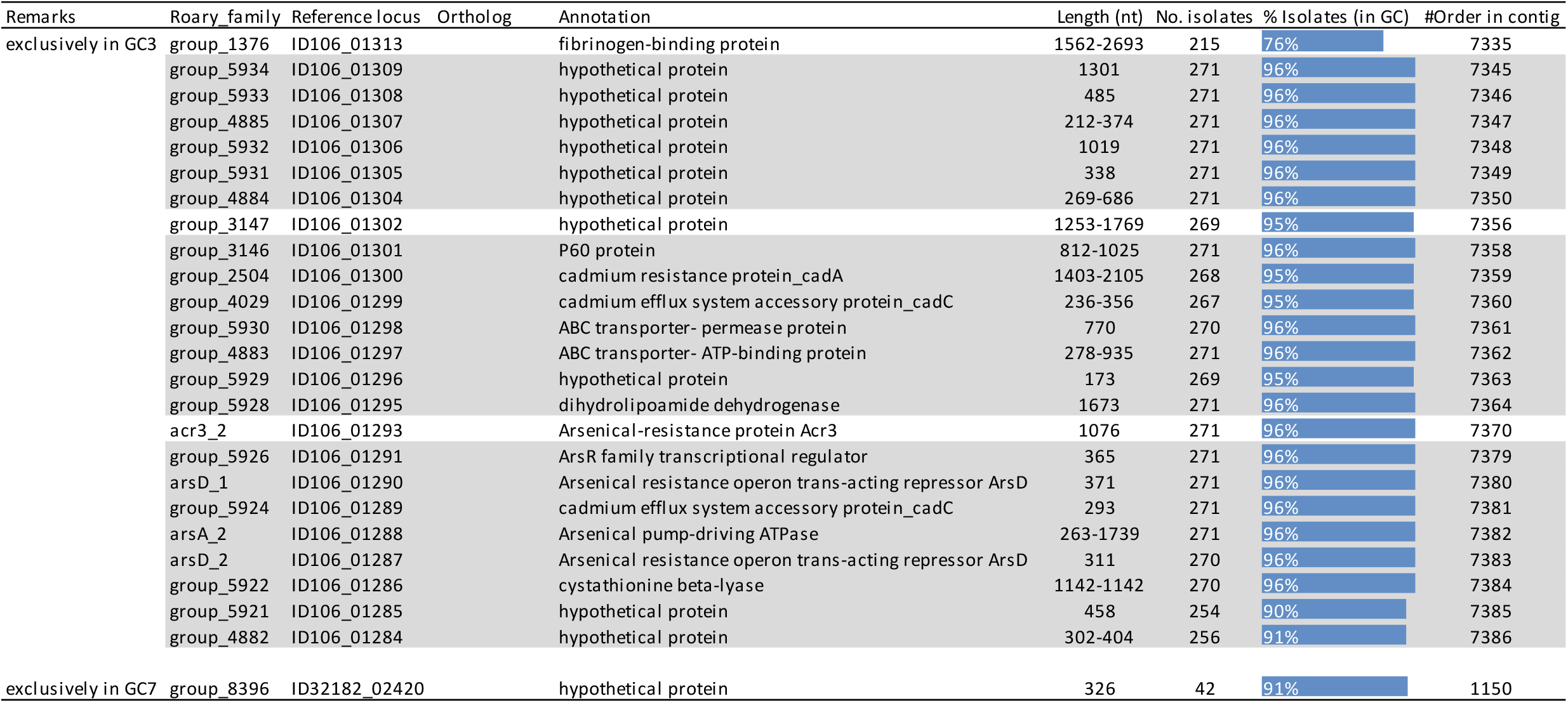
SL1 genetic clades-specific genes present in at least 50% of isolates. Clades-specific genes were only found in GC3 and GC7. Gray shadows highlight genes within the same genomic context.

| Remarks | Roary_family | Reference locus | Ortholog | Annotation | Length (nt) | No. isolates | % Isolates (in GC) | #Order in contig |
| --- | --- | --- | --- | --- | --- | --- | --- | --- |
| exclusively in GC3 | group_1376 | ID106_01313 |  | fibrinogen-binding protein | 1562-2693 | 215 | 76% | 7335 |
|  | group_5934 | ID106_01309 |  | hypothetical protein | 1301 | 271 | 96% | 7345 |
|  | group_5933 | ID106_01308 |  | hypothetical protein | 485 | 271 | 96% | 7346 |
|  | group_4885 | ID106_01307 |  | hypothetical protein | 212-374 | 271 | 96% | 7347 |
|  | group_5932 | ID106_01306 |  | hypothetical protein | 1019 | 271 | 96% | 7348 |
|  | group_5931 | ID106_01305 |  | hypothetical protein | 338 | 271 | 96% | 7349 |
|  | group_4884 | ID106_01304 |  | hypothetical protein | 269-686 | 271 | 96% | 7350 |
|  | group_3147 | ID106_01302 |  | hypothetical protein | 1253-1769 | 269 | 95% | 7356 |
|  | group_3146 | ID106_01301 |  | P60 protein | 812-1025 | 271 | 96% | 7358 |
|  | group_2504 | ID106_01300 |  | cadmium resistance protein_cadA | 1403-2105 | 268 | 95% | 7359 |
|  | group_4029 | ID106_01299 |  | cadmium efflux system accessory protein_cadC | 236-356 | 267 | 95% | 7360 |
|  | group_5930 | ID106_01298 |  | ABC transporter- permease protein | 770 | 270 | 96% | 7361 |
|  | group_4883 | ID106_01297 |  | ABC transporter- ATP-binding protein | 278-935 | 271 | 96% | 7362 |
|  | group_5929 | ID106_01296 |  | hypothetical protein | 173 | 269 | 95% | 7363 |
|  | group_5928 | ID106_01295 |  | dihydrolipoamide dehydrogenase | 1673 | 271 | 96% | 7364 |
|  | acr3_2 | ID106_01293 |  | Arsenical-resistance protein Acr3 | 1076 | 271 | 96% | 7370 |
|  | group_5926 | ID106_01291 |  | ArsR family transcriptional regulator | 365 | 271 | 96% | 7379 |
|  | arsD_1 | ID106_01290 |  | Arsenical resistance operon trans-acting repressor ArsD | 371 | 271 | 96% | 7380 |
|  | group_5924 | ID106_01289 |  | cadmium efflux system accessory protein_cadC | 293 | 271 | 96% | 7381 |
|  | arsA_2 | ID106_01288 |  | Arsenical pump-driving ATPase | 263-1739 | 271 | 96% | 7382 |
|  | arsD_2 | ID106_01287 |  | Arsenical resistance operon trans-acting repressor ArsD | 311 | 270 | 96% | 7383 |
|  | group_5922 | ID106_01286 |  | cystathionine beta-lyase | 1142-1142 | 270 | 96% | 7384 |
|  | group_5921 | ID106_01285 |  | hypothetical protein | 458 | 254 | 90% | 7385 |
|  | group_4882 | ID106_01284 |  | hypothetical protein | 302-404 | 256 | 91% | 7386 |
| exclusively in GC7 | group_8396 | ID32182_02420 |  | hypothetical protein | 326 | 42 | 91% | 1150 |

**Table S5.** Human-associated significant loci, as determined using treeWAS, with a significance threshold of *p*<10^−5^.

| Gene | Annotation | treeWAS score | Association type | G1P1 | G0P0 | G1P0 | G0P1 |
| --- | --- | --- | --- | --- | --- | --- | --- |
| group_1361 | hypothetical protein | 24 | positive | 1303 | 88 | 480 | 150 |
| group_2465 | valyl-tRNA synthetase_valS | 22 | positive | 1300 | 89 | 479 | 153 |
| group_1038 | N-acetylmuramoyl-L-alanine amidase | 26 | positive | 1265 | 96 | 472 | 188 |
| group_619 | hypothetical protein | 23 | positive | 1175 | 138 | 430 | 278 |
| group_10387 | tRNA-Glu(ttc) | 25 | positive | 1041 | 181 | 387 | 412 |
| group_497 | hypothetical protein | 24 | positive | 1029 | 190 | 378 | 424 |
| group_1926 | transcriptional regulator | 25 | positive | 1024 | 193 | 375 | 429 |
| group_706 | hypothetical protein | 25 | positive | 1020 | 191 | 377 | 433 |
| group_1527 | hypothetical protein | 24 | positive | 1007 | 186 | 382 | 446 |
| group_209 | hypothetical protein | 24 | positive | 866 | 254 | 314 | 587 |
| group_6398 | tRNA-Val(tac) | -25 | negative | 611 | 246 | 322 | 842 |
| group_10476 | 5S ribosomal RNA | -31 | negative | 520 | 253 | 315 | 933 |
| group_10390 | tRNA-Glu(ttc) | -33 | negative | 532 | 276 | 292 | 921 |
| group_10432 | tRNA-Lys(ttt) | -35 | negative | 499 | 289 | 279 | 954 |
| group_10162 | tRNA-Asn(gtt) | -29 | negative | 461 | 309 | 259 | 992 |
| group_10094 | hypothetical protein | -28 | negative | 492 | 362 | 206 | 961 |
| group_10662 | hypothetical protein | -30 | negative | 479 | 358 | 210 | 974 |
| group_1927 | transcriptional regulator | -25 | negative | 430 | 375 | 193 | 1023 |
| group_499 | hypothetical protein | -23 | negative | 404 | 397 | 171 | 1049 |
| group_10488 | 5S ribosomal RNA | -41 | negative | 282 | 399 | 169 | 1171 |
| group_4211 | 5S ribosomal RNA (partial) | -37 | negative | 227 | 398 | 170 | 1226 |
| group_60 | putative lipoprotein | -27 | negative | 188 | 476 | 92 | 1265 |
| group_533 | hypothetical protein | -22 | negative | 74 | 508 | 60 | 1379 |
| group_6404 | hypothetical protein | -22 | negative | 36 | 518 | 50 | 1417 |

G, genome; P, phenotype; 0 absent, 1 present.

### 2. Supplementary figures

**Figure S1.**
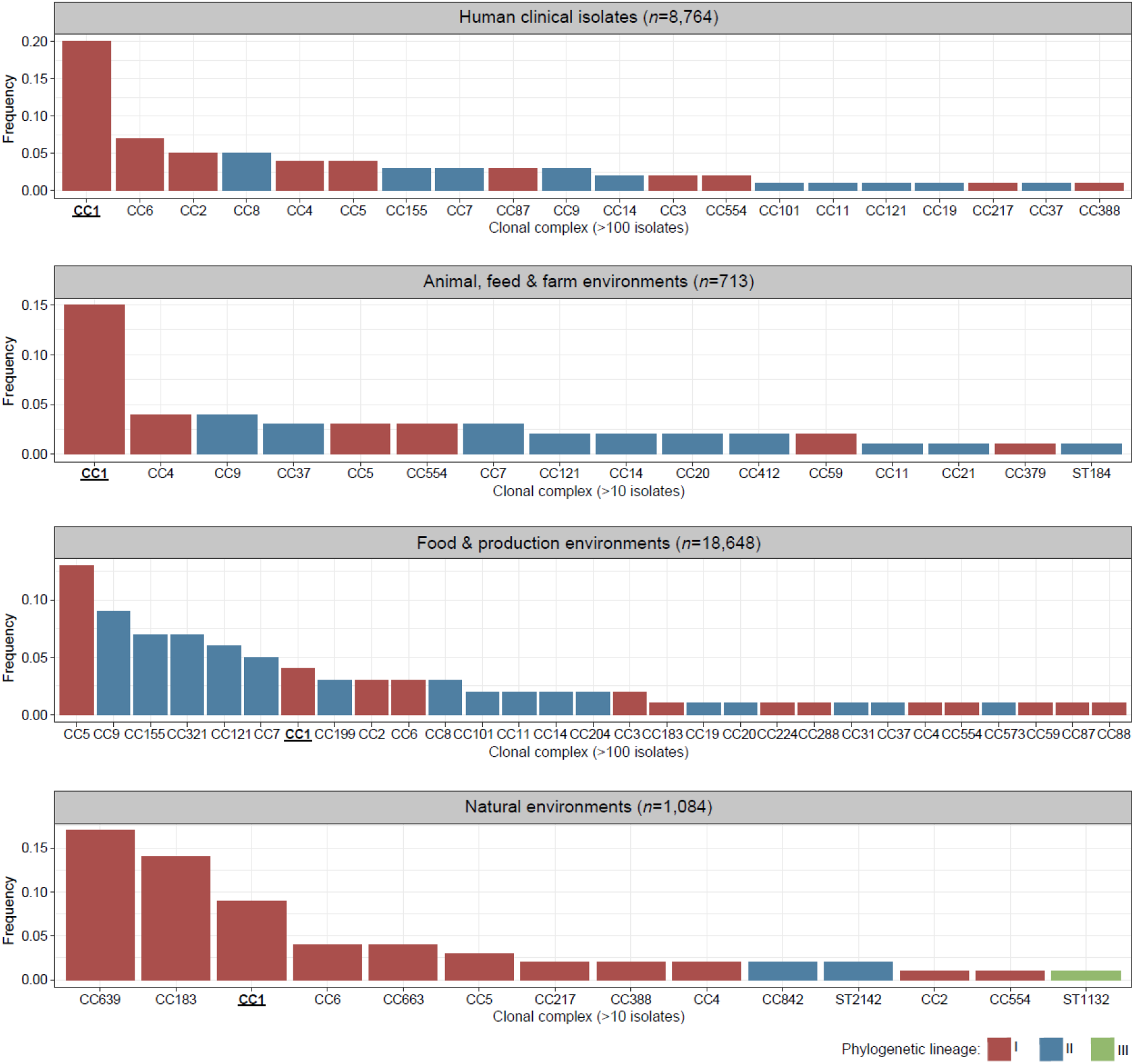
Frequency of most prevalent clonal complexes among different environments. Data collected based on 29,349 *L. monocytogenes* genomes with associated source metadata available on NCBI Sequence Read Archive (as of October 23^rd^, 2020). MLST typing was performed from reads using the srst2 v.0.1.5 software (http://katholt.github.io/srst2) and the BIGSdb-*Lm* profiles database (https://bigsdb.pasteur.fr/listeria).

**Figure S2.**
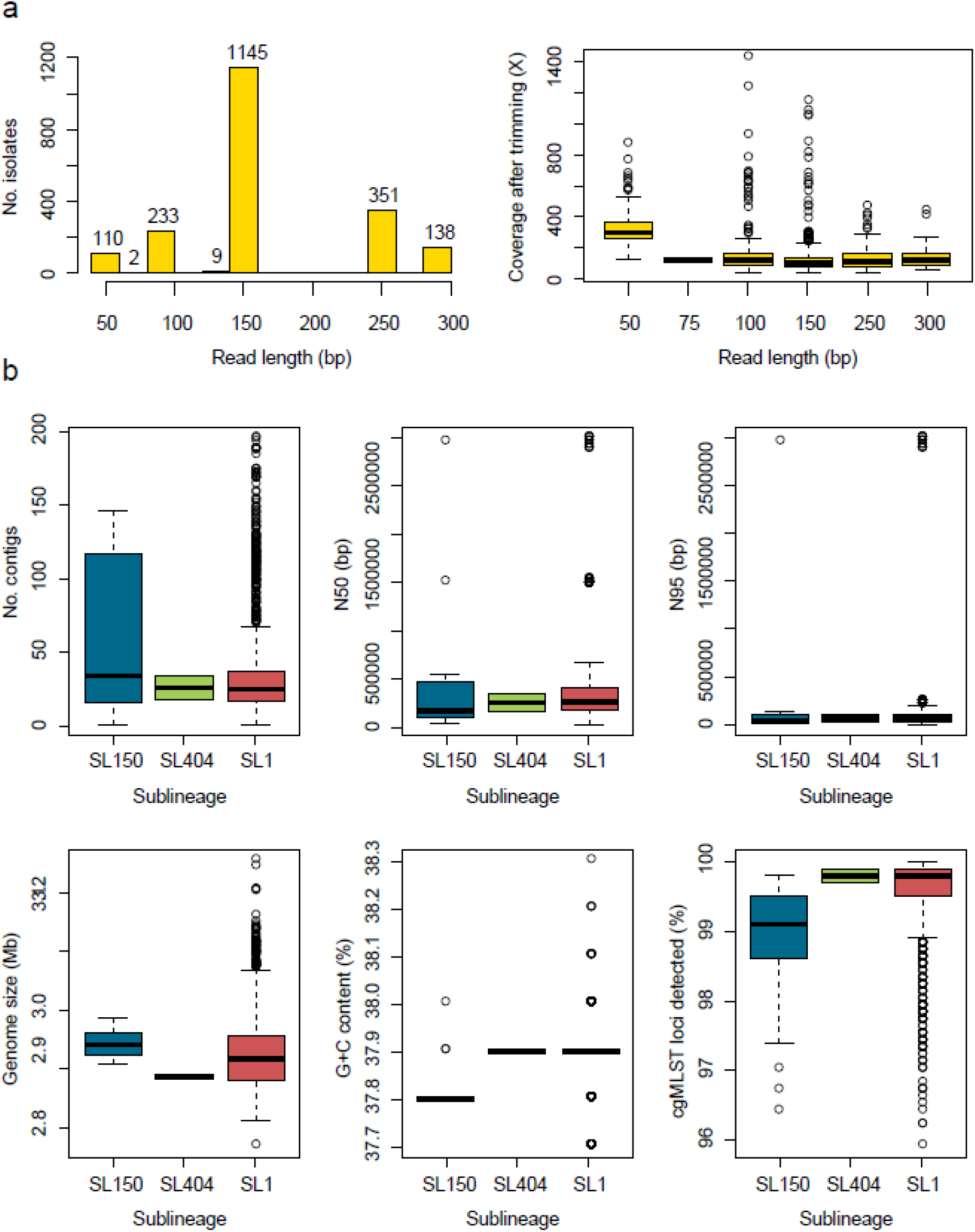
Genome metrics of isolates included in this study. a) Distribution of isolates per sequence read length (left) and distribution of sequencing coverages after reads quality trimming (right). b) Assembly metrics per CC1 sublineages, based on the number of contigs, N50 and N95 contig lengths, genome size, G+C content and cgMLST loci detected.

**Figure S3.**
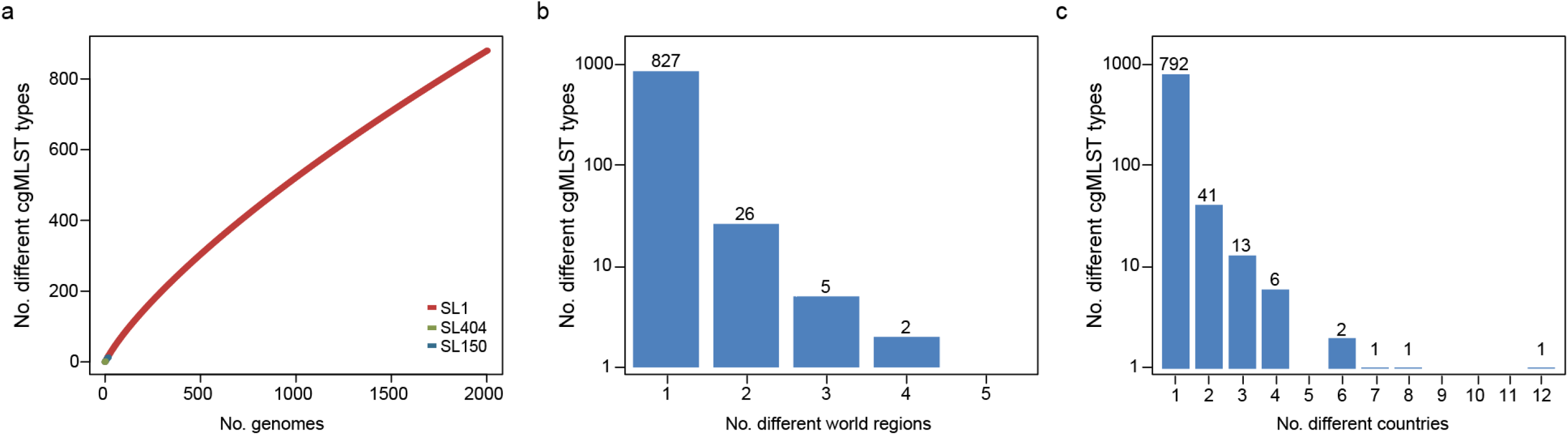
Core genome multilocus sequence typing (cgMLST) analyses. a) Rarefaction analysis of cgMLST types sampled per sublineage. b) Number of SL1 cgMLST types per number of different world regions in which they were observed (*n*=860 types with world region information). c) Number of SL1 cgMLST types per number of different countries in which they were observed (*n*=857 types with country information).

**Figure S4.**
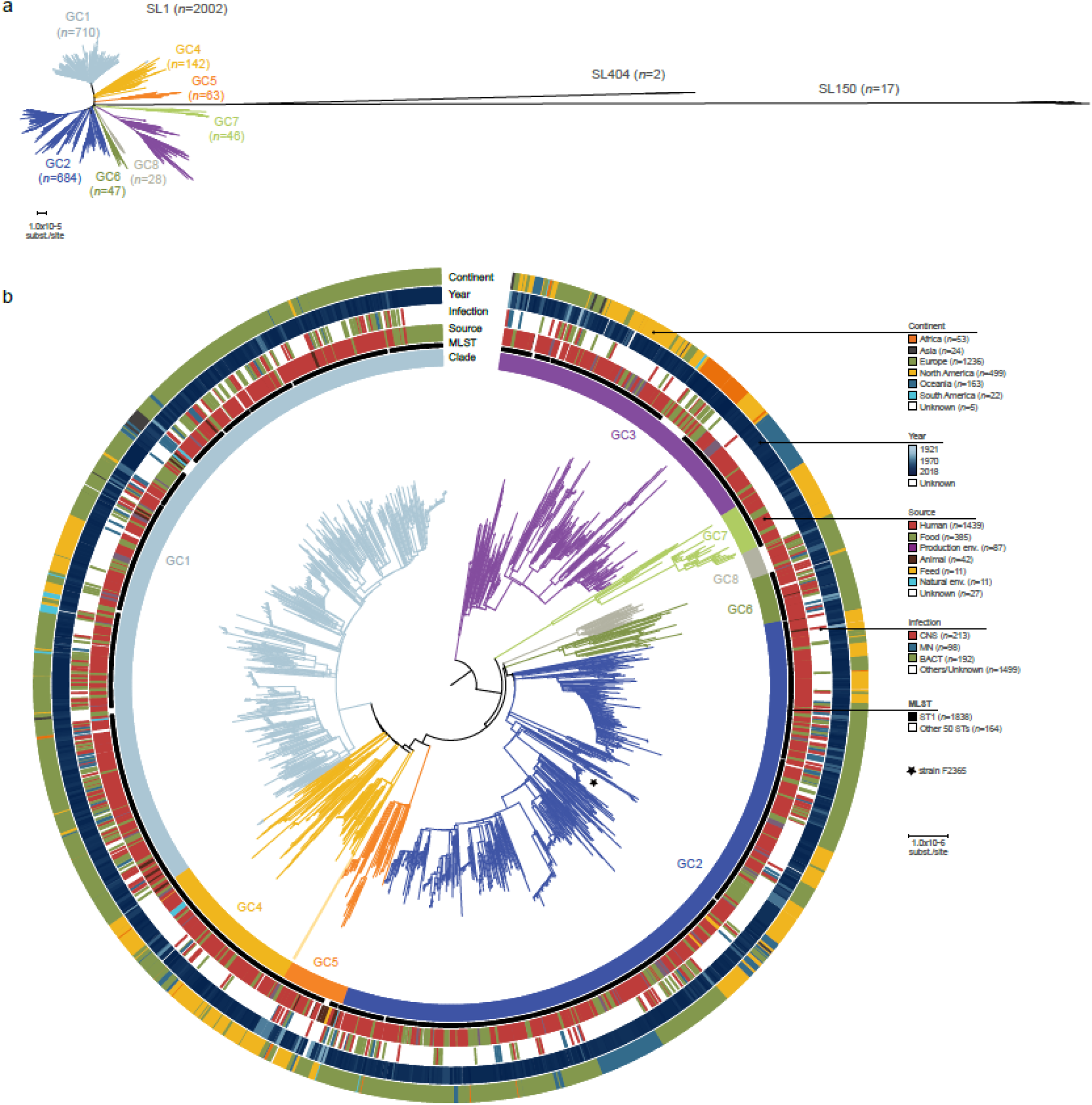
Phylogenetic analysis based on whole genome SNP analyses. a) Unrooted maximum-likelihood phylogeny (GTR+F+G4 model, 1000 ultra-fast bootstraps, using IQ-Tree^23,26^) of 2,021 CC1 genomes based on the recombination-purged whole genome SNP alignment of 2.28 Mb. b) Midpoint rooted maximum-likelihood phylogenetic tree of 2,002 SL1 genomes based on based on the recombination-purged whole genome SNP alignment of 2.28 Mb. The four external rings indicate the world region, year, type of infection and source type, respectively. The two inner rings indicate ST1 isolates and the 8 SL1 genetic clades identified in this study, respectively. The black star highlights the phylogenetic placement of isolate F2365 (accession no. NC_002973.6), used as reference in whole genome read mapping.

**Figure S5.**
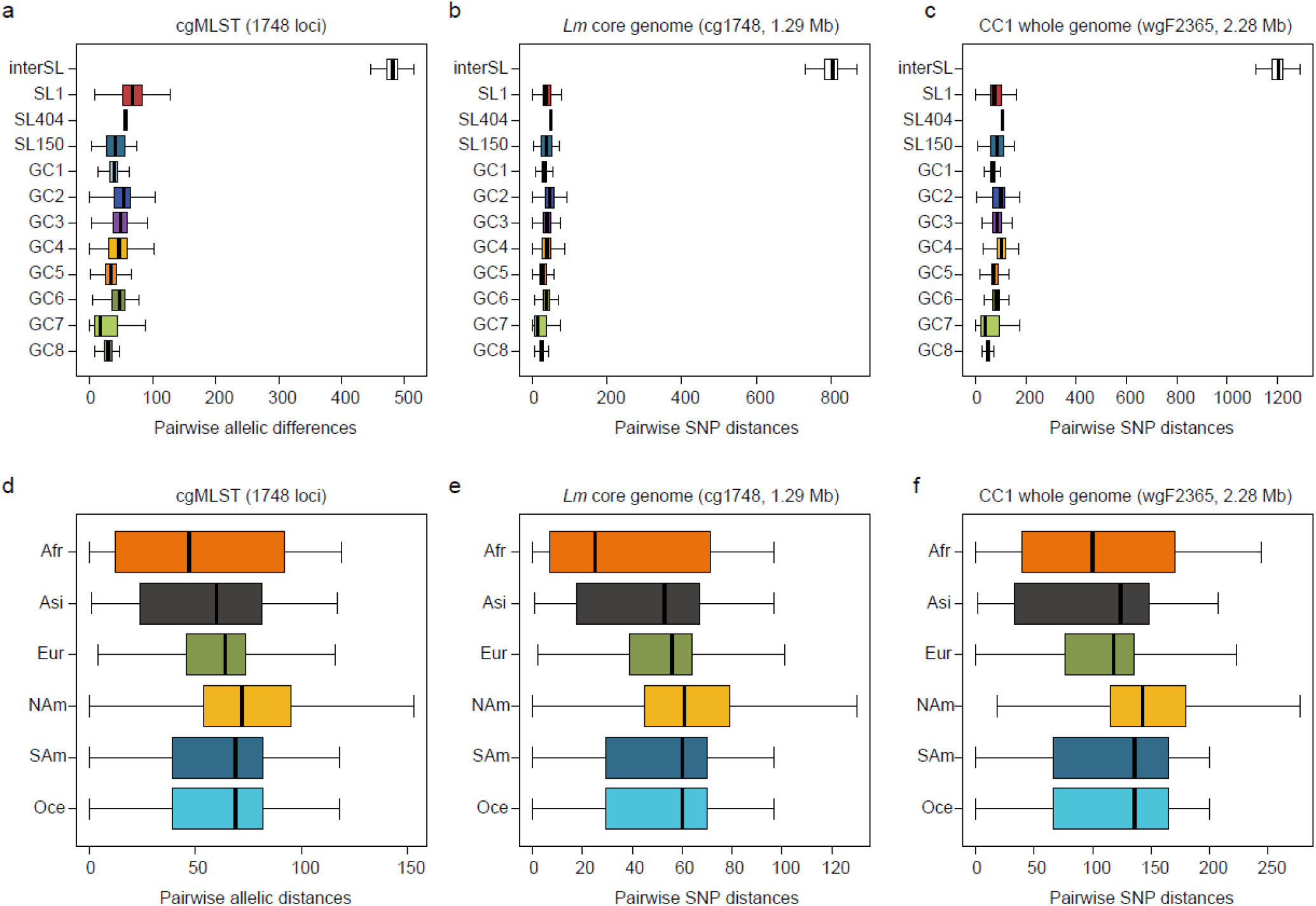
Genetic diversity among *Lm*-CC1 isolates. Pairwise isolate distances within CC1 phylogroups (top) and world regions (bottom): a,d) pairwise cgMLST allelic distances; b,e) pairwise SNP distances in recombination-purged *Lm* core genome alignment and c,f) recombination-purged CC1 whole genome alignment. Uncalled alleles, Ns and gap alignment positions were ignored in pairwise comparisons. Each box denotes the 25% and 75% quartiles and lines represent the medians. Inter-SL, inter CC1 sublineages; GC#, within SL1 genetic clades; Afr, Africa; Asi, Asia; Eur, Europe; NAm, North America; Sam, South America; Oce, Oceania.

**Figure S6.**
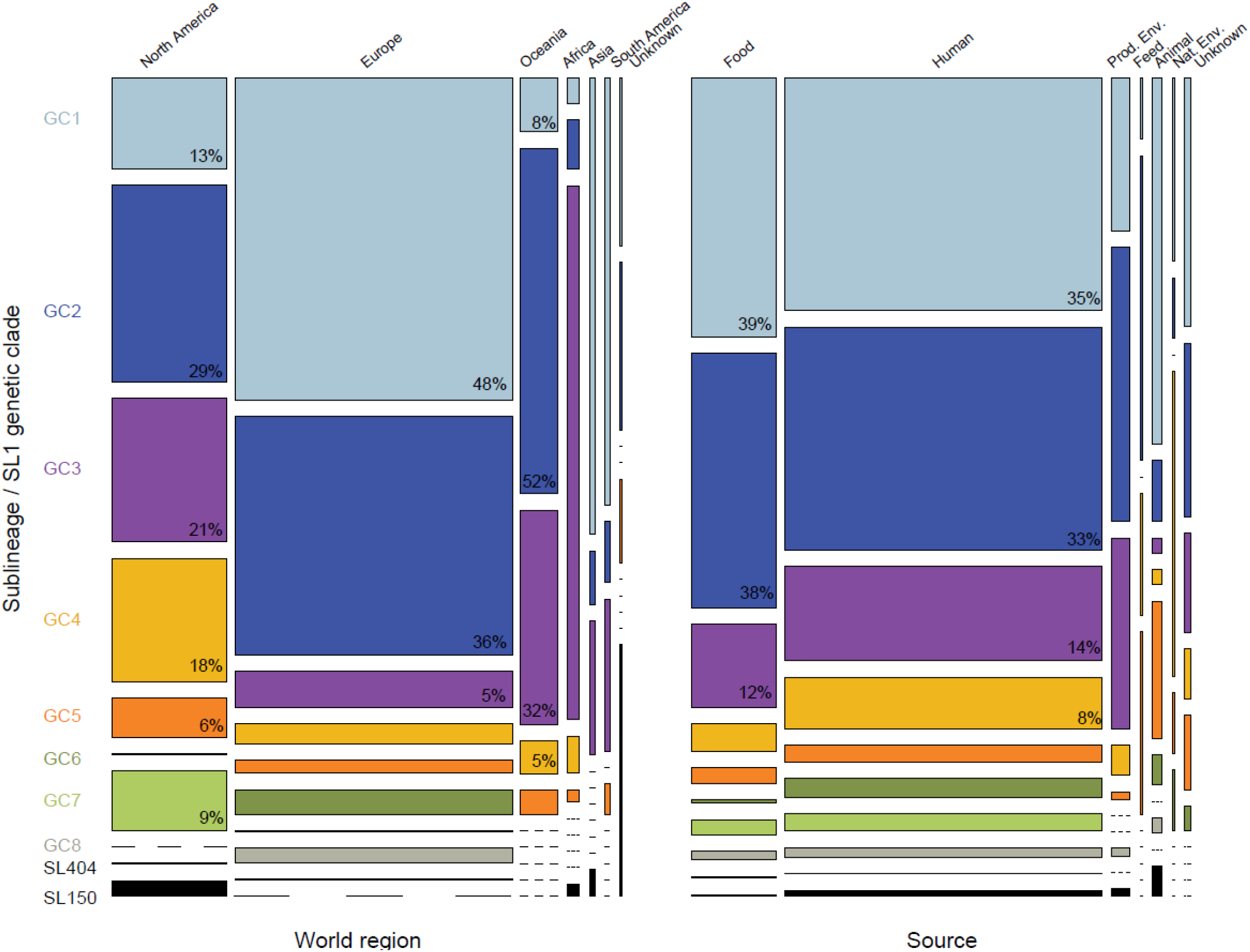
Distribution of *Lm*-CC1 isolates per clade, world regions and source types (*N*=2,021).

**Figure S7.**
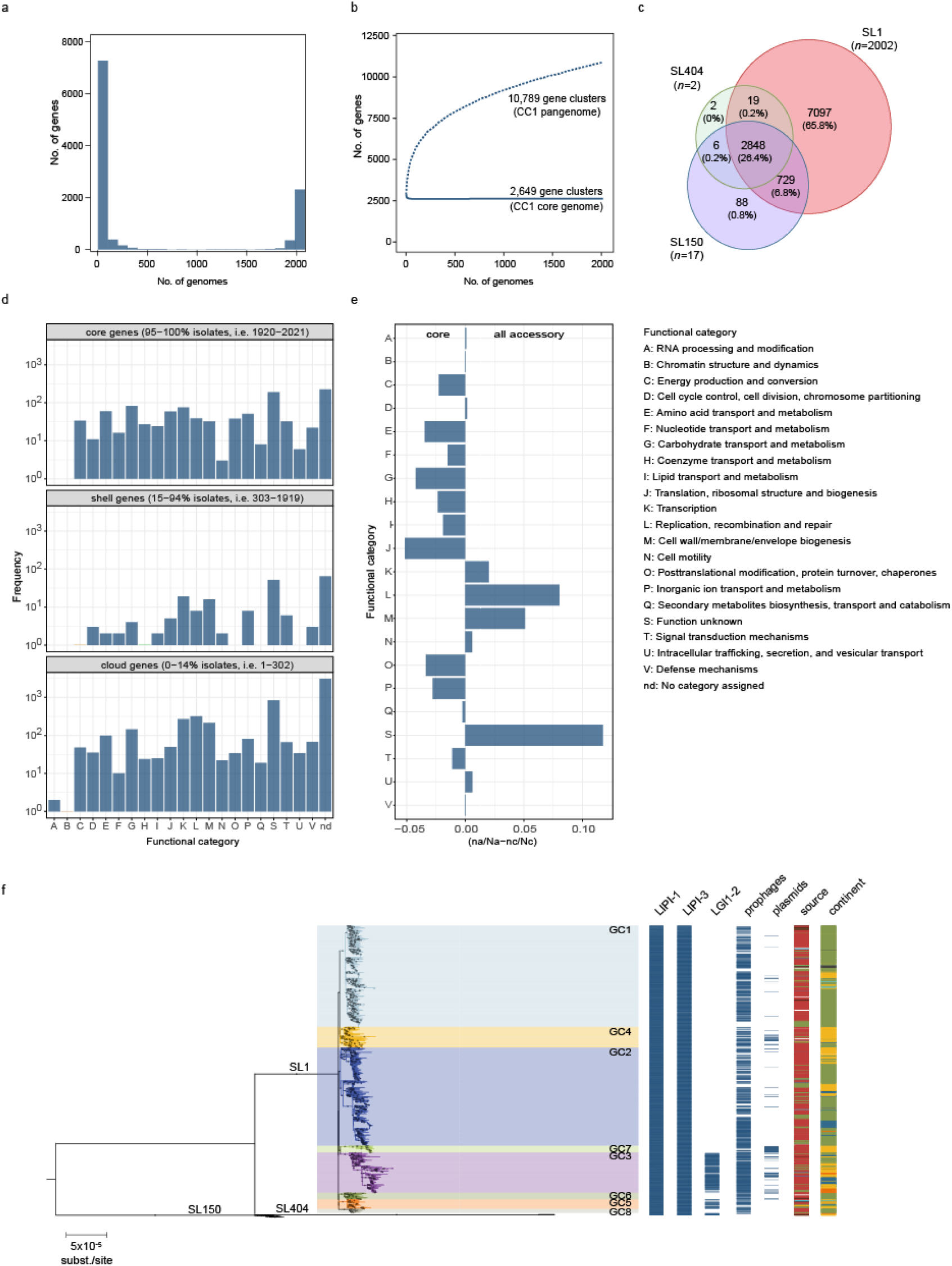
CC1 pangenome analysis. a) Frequency of sampled gene families. b) Pan- and core gene families sampled. c) Venn diagram showing the number of gene families present in at least 1 sublineage member. d) Distribution of the functional categories of the clusters of orthologous genes across the CC1 pangenome. e) Differential proportion of each assigned COG category in core vs accessory genome, calculated as the difference between the ratio of each category (*n*) and the total number of hits (*N*) among each gene pool set, as in (*n*_accessory_/*N*_accessory_-*n*_core_/*N*_core_). f) Distribution of *Listeria* genomic islands, prophages and plasmids and across CC1 phylogeny. The midpoint rooted maximum-likelihood phylogenetic tree (GTR+F+G4 model, 1000 ultra-fast bootstraps) was inferred from the 1.29 Mb recombination-purged core genome alignment of 2,021 CC1 genomes. Sources, continents and SL1 clades are colored according to the color codes shown in Figure S4.

**Figure S8.**
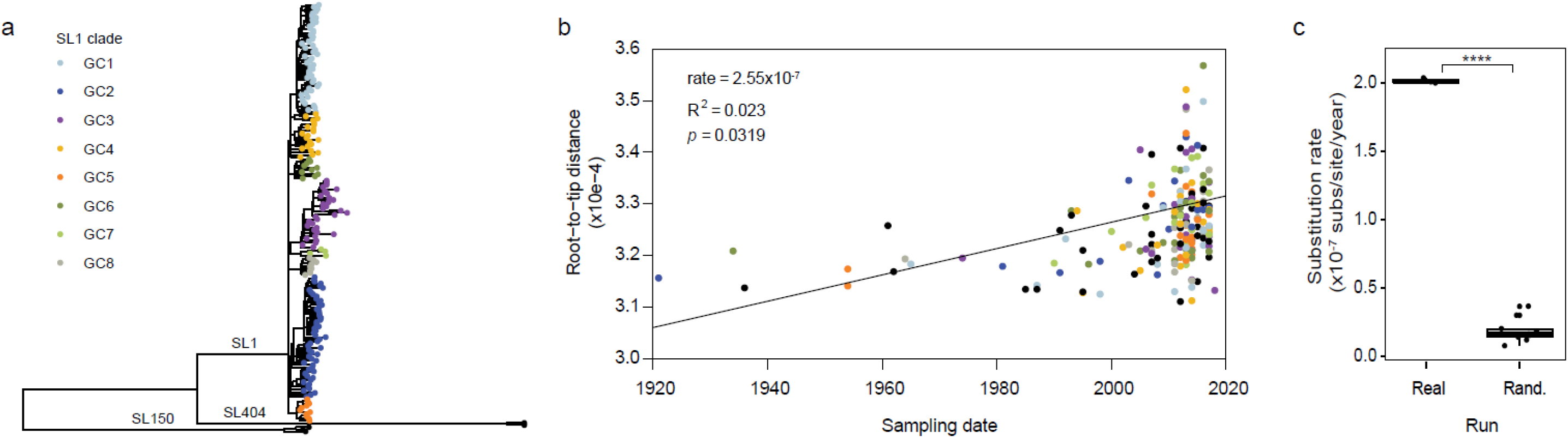
Temporal analyses on a representative dataset of 200 isolates. a) Maximum likelihood (GTR+F+G4) phylogeny of the representative 200 isolates selected randomly across the CC1 phylogeny. Tips are colored by sublineage and SL1 genetic clades as indicated in the legend. b) Regression analyses showing the root-to-tip genetic distance against sampling date (year). Statistical significance was assessed using the F-test. c) Bayesian molecular clock estimations in real and randomized tip dates (controls). Estimations based on real data were run in triplicates, whereas estimations based on randomized tip datasets were run in 10 replicates. Stars denote statistical significance of *p*<0.0001, assessed using t-test.

**Figure S9.**
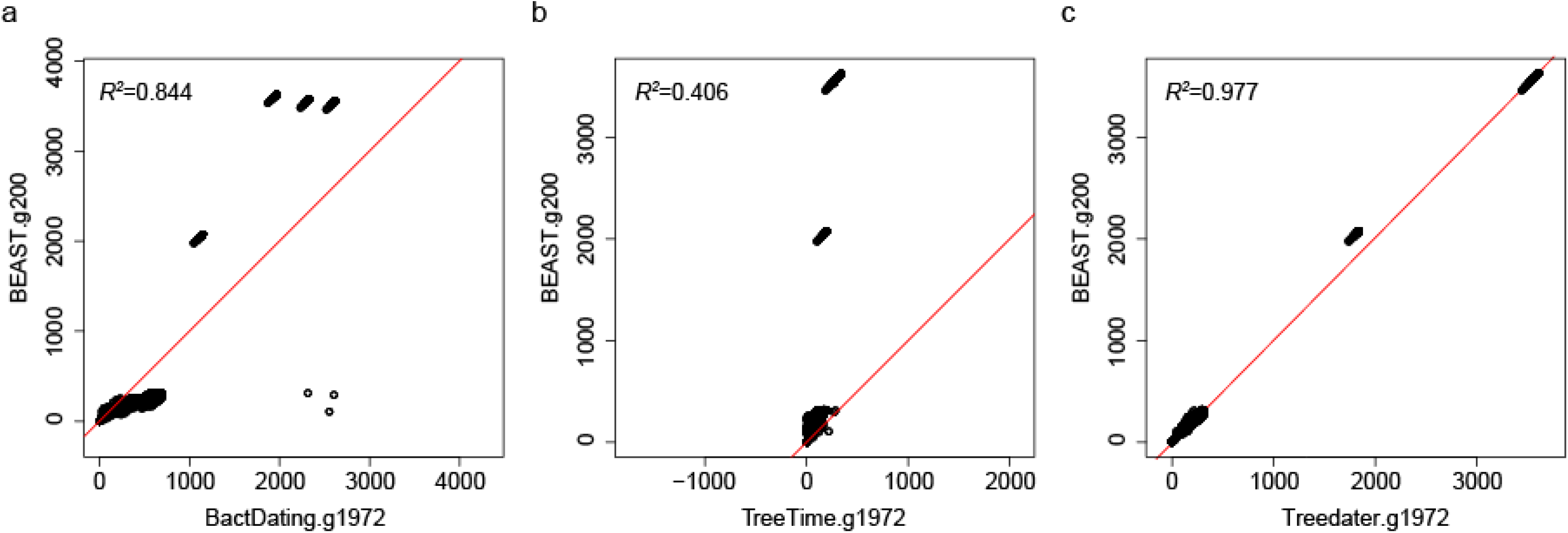
Benchmarking of dating methods. Cophenetic distances between isolates dated with BEAST and alternative large-scale methods: a) BactDating v.1.0.1, b) Treetime v.0.5.2 and c) Treedater v.0.3.0, using the CC1 estimated rate of 1.954×10^−7^ ± 2.0152×10^−8^ substitutions/site/year obtained with BEAST. “g200” and “g1972” refer to the number of CC1 genomes used in each analyses (*n*=200 and *n*=1,972, respectively). Red lines denote perfect positive correlation coefficients (*R^2^*=1).

**Figure S10.**
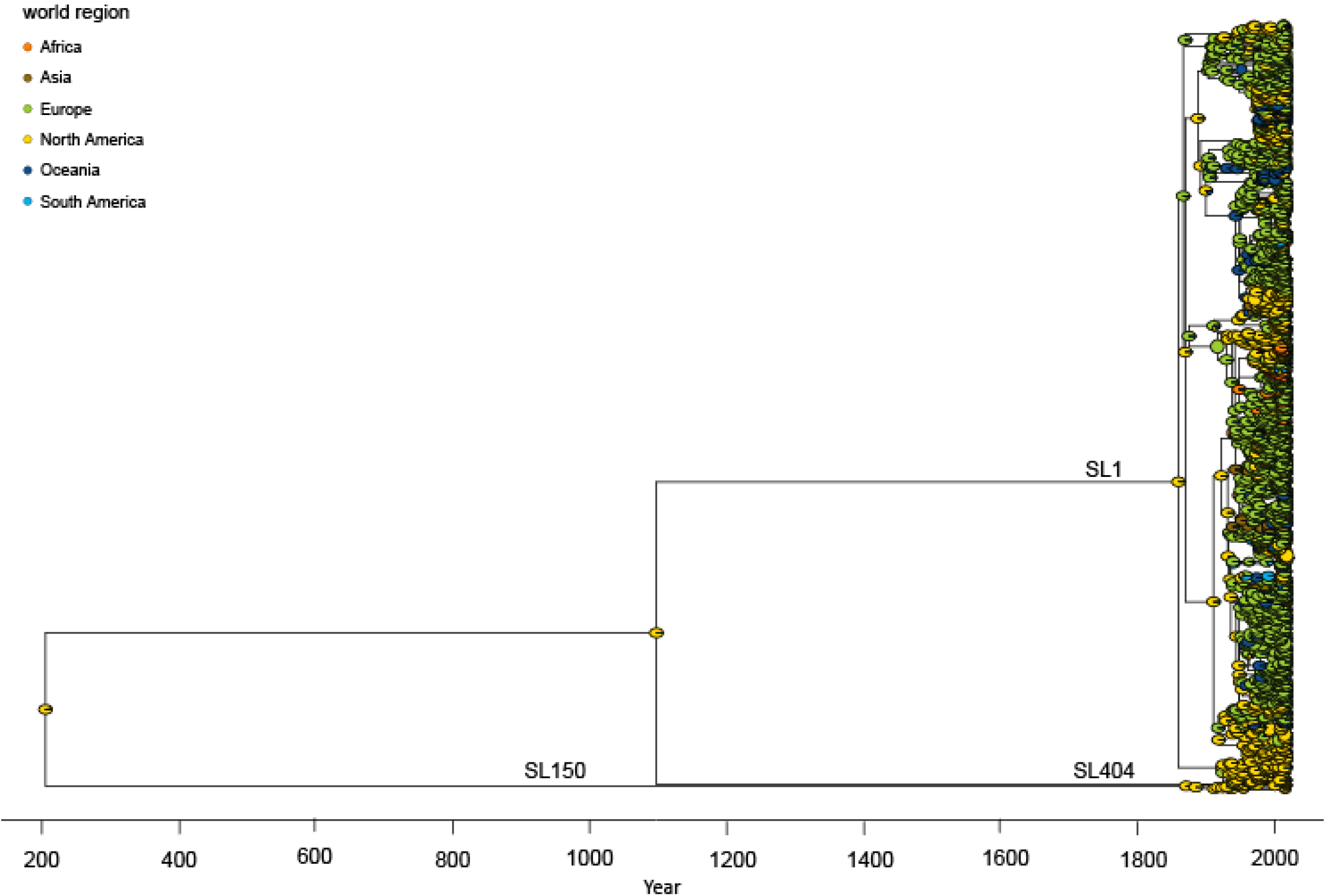
Phylogeography inference of *Lm*-CC1 based on 1972 dated genomes. Pies at the nodes represent the probability of ancestral geographical locations, estimate using PastML using the MPPA method with an F81-like model. The detailed view of SL1 can be found in Figure 3.

**Figure S11.**
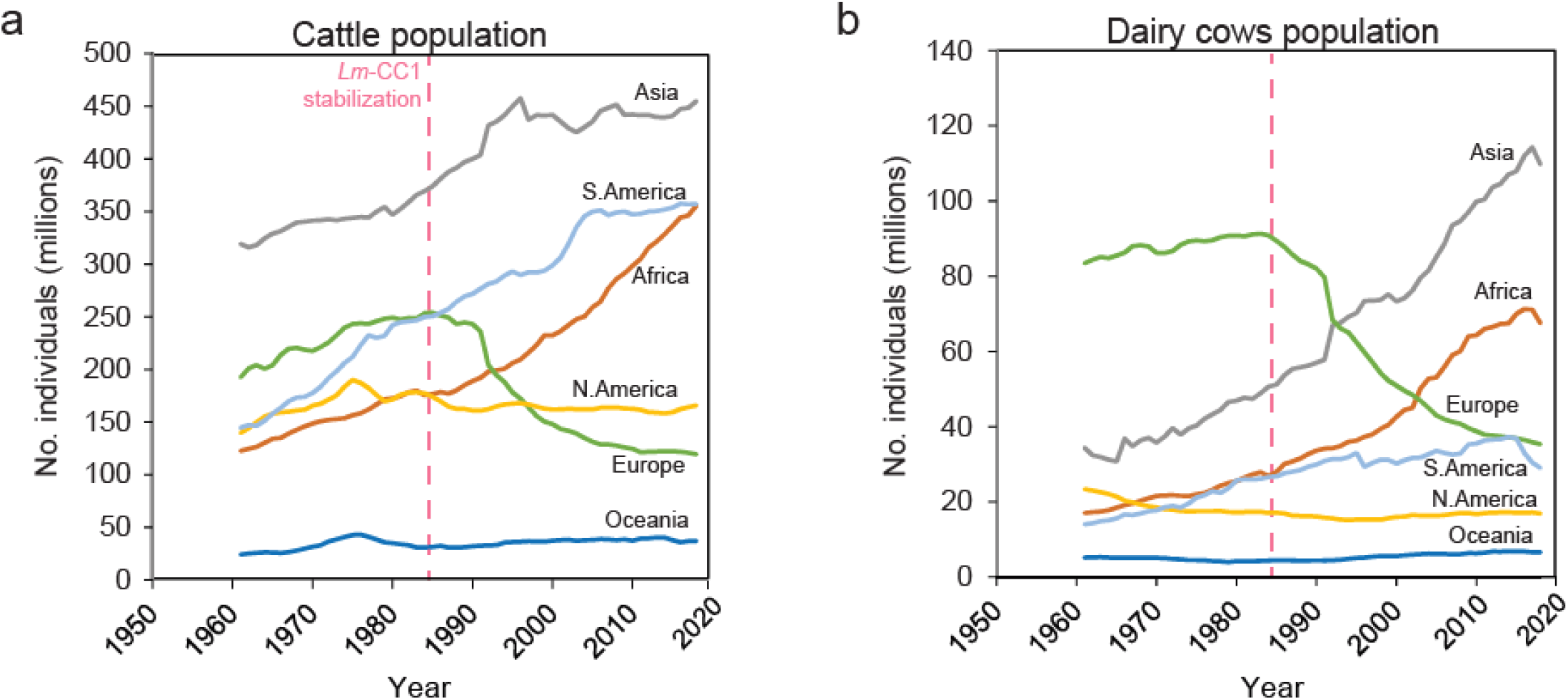
Cattle demographics. a) Cattle population per world region; b) Dairy cows per world region. Data available for 1961-2018; source: Food and Agriculture Organization of the United Nations; www.fao.org/faostat). Vertical dashed bars mark the estimated date of the stabilization *of Lm*-CC1 population size.

**Figure S12.**
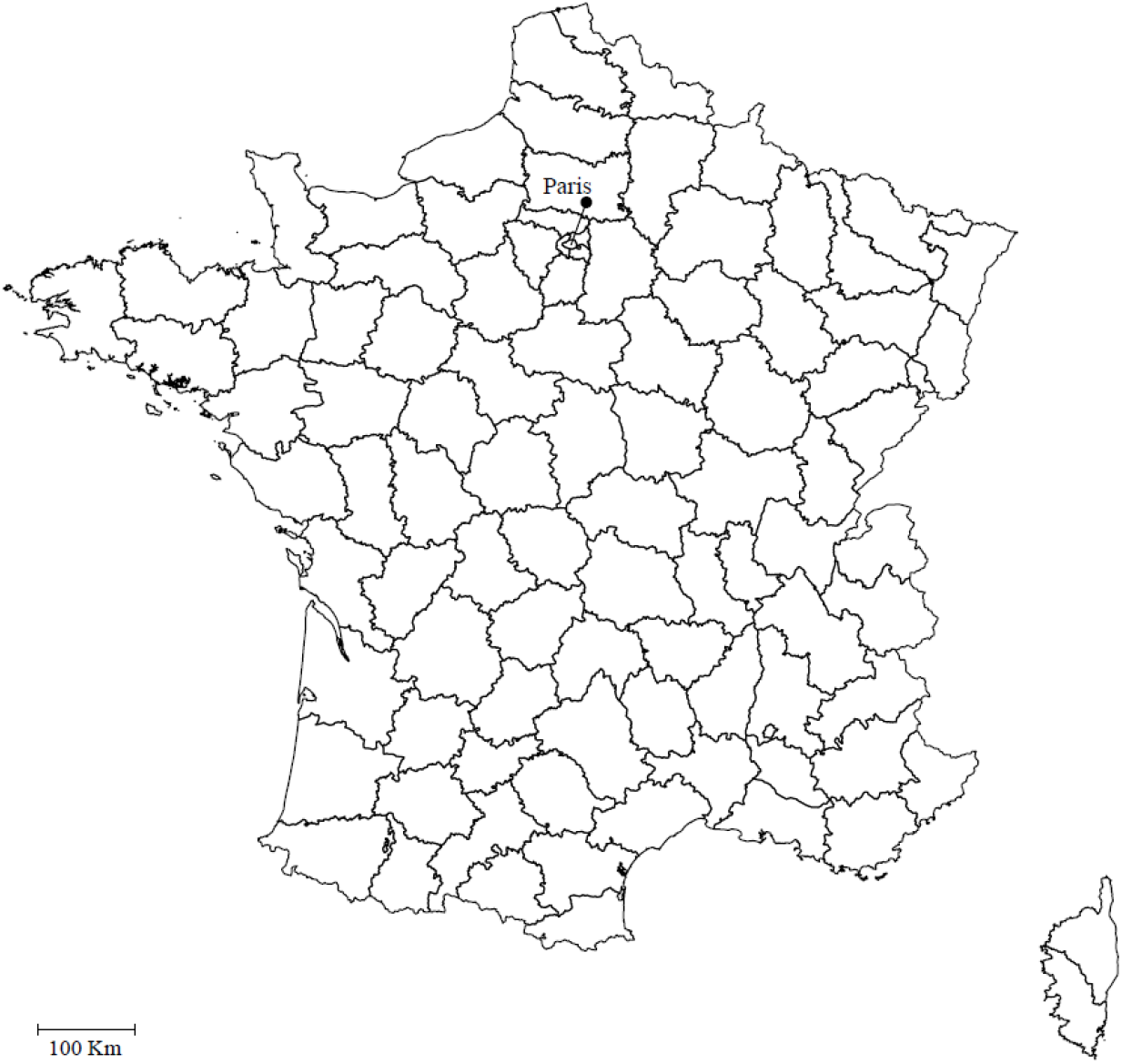
French administrative Departments(*départements*). Source: Global Administrative Areas, gadm.org.

**Figure S13.**
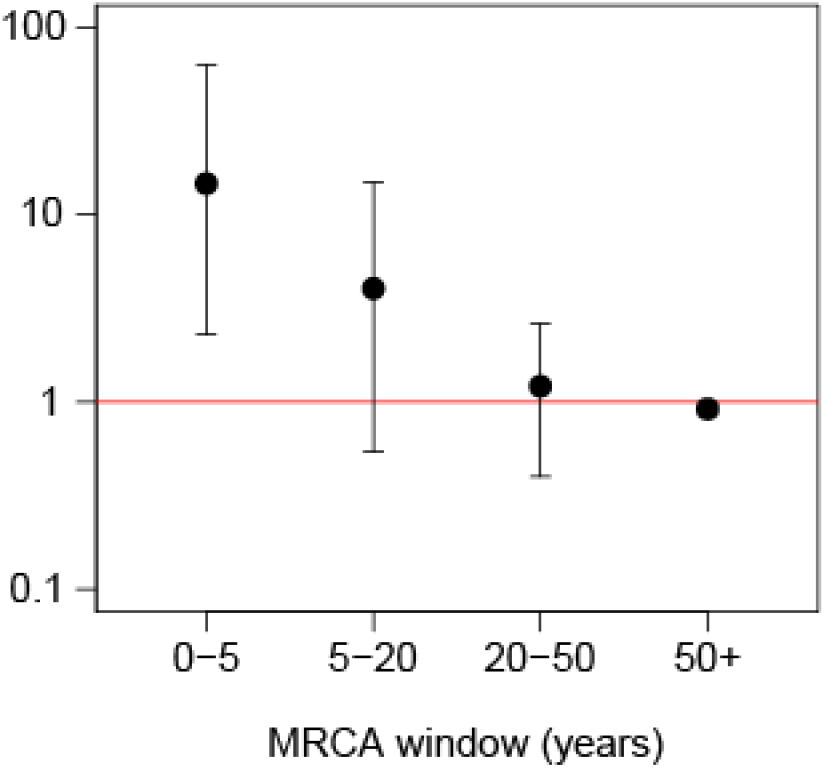
SL1 transmission dynamics within country. (France). Relative risk for a pair of isolates to have a MRCA within a defined period when coming from the same Department in France *versus* different ones.

## References

1. Swaminathan, B. & Gerner-Smidt, P. The epidemiology of human listeriosis. Microbes Infect. 9, 1236–1243 (2007).

2. Charlier, C. et al. Clinical features and prognostic factors of listeriosis: the MONALISA national prospective cohort study. Lancet Infect. Dis. 17, 510–519 (2017).

3. Orsi, R. H., Bakker, H. C. de. & Wiedmann, M. *Listeria monocytogenes* lineages: genomics, evolution, ecology, and phenotypic characteristics. Int. J. Med. Microbiol. 301, 79–96 (2011).

4. Ragon, M. et al. A new perspective on *Listeria monocytogenes* evolution. PLoS Pathog. 4, e1000146 (2008).

5. Cantinelli, T. et al. ‘Epidemic clones’ of *Listeria monocytogenes* are widespread and ancient clonal groups. J. Clin. Microbiol. 51, 3770–3779 (2013).

6. Chenal-Francisque, V. et al. Worldwide distribution of major clones of *Listeria monocytogenes*. Emerg. Infect. Dis. 17, 1110–1112 (2011).

7. Dumont, J. & Cotoni, L. Bacille semblable au bacielle du Rouget du porc rencontré dans le liquide céphalo-rachidien d’un méningitique. Ann. Inst. Pasteur (Paris). 35, 625–633 (1921).

8. Hyden, P. et al. Draft genome sequence of a 94-year-old *Listeria monocytogenes* isolate, SLCC208. Genome Announc. 4, e01572–15 (2016).

9. Maury, M. et al. Uncovering *Listeria monocytogenes* hypervirulence by harnessing its biodiversity. Nat. Genet. 48, 308–313 (2016).

10. Kwong, J. C. et al. Prospective whole genome sequencing enhances national surveillance of *Listeria monocytogenes*. J. Clin. Microbiol. 54, JCM.02344–15 (2015).

11. Bertrand, S. et al. Diversity of *Listeria monocytogenes* strains of clinical and food chain origins in Belgium between 1985 and 2014. PLoS One 11, e0164283 (2016).

12. Toledo, V. et al. Genomic diversity of *Listeria monocytogenes* isolated from clinical and non-clinical samples in Chile. Genes (Basel). 9, 396 (2018).

13. Hilliard, A. et al. Genomic characterization of *Listeria monocytogenes* isolates associated with clinical listeriosis and the food production environment in Ireland. Genes (Basel). 9, 171 (2018).

14. Scaltriti, E. et al. Population Structure of *Listeria monocytogenes* in Emilia-Romagna (Italy) and implications on whole genome sequencing surveillance of listeriosis. Front. Public Heal. 8, (2020).

15. Schlech, W. F. et al. Epidemic listeriosis — evidence for transmission by food. N. Engl. J. Med. 308, 203–206 (1983).

16. Maury, M. M. et al. Hypervirulent *Listeria monocytogenes* clones’ adaption to mammalian gut accounts for their association with dairy products. Nat. Commun. 10, 2488 (2019).

17. Painset, A. et al. Liseq – Whole-genome sequencing of a cross-sectional survey of *Listeria monocytogenes* in ready-to-eat foods and human clinical cases in Europe. Microb. Genomics 5, e000257 (2019).

18. Félix, B. et al. Population genetic structure of *Listeria monocytogenes* strains isolated from the pig and pork production chain in France. Front. Microbiol. 9, (2018).

19. Dalton, C. B. et al. An outbreak of gastroenteritis and fever due to *Listeria monocytogenes* in milk. N. Engl. J. Med. 336, 100–5 (1997).

20. Costard, S., Espejo, L., Groenendaal, H. & Zagmutt, F. J. Outbreak-related disease burden associated with consumption of unpasteurized cow’s milk and cheese, United States, 2009–2014. Emerg. Infect. Dis. 23, 957–964 (2017).

21. Filipello, V. et al. Attribution of *Listeria monocytogenes* human infections to food and animal sources in Northern Italy. Food Microbiol. 89, (2020).

22. Dreyer, M. et al. *Listeria monocytogenes* sequence type 1 is predominant in ruminant rhombencephalitis. Sci. Rep. 6, 36419 (2016).

23. Papić, B., Pate, M., Félix, B. & Kušar, D. Genetic diversity of *Listeria monocytogenes* strains in ruminant abortion and rhombencephalitis cases in comparison with the natural environment. BMC Microbiol. 19, 299 (2019).

24. Garcia-Garcera, M. et al. *Listeria monocytogenes* faecal carriage is common and driven by microbiota. bioRkiv (2020).

25. Nightingale, K. K. et al. Ecology and transmission of *Listeria monocytogenes* infecting ruminants and in the farm environment. Appl. Environ. Microbiol. 70, 4458–4467 (2004).

26. Esteban, J. I., Oporto, B., Aduriz, G., Juste, R. A. & Hurtado, A. Faecal shedding and strain diversity of *Listeria monocytogenes* in healthy ruminants and swine in Northern Spain. BMC Vet. Res. 5, (2009).

27. Lyautey, E. et al. Characteristics and frequency of detection of fecal *Listeria monocytogenes* shed by livestock, wildlife, and humans. Can. J. Microbiol. 53, 1158–1167 (2007).

28. Borucki, M. K. et al. Genetic diversity of *Listeria monocytogenes* strains from a high-prevalence dairy farm. Appl. Environ. Microbiol. 71, 5893–5899 (2005).

29. Jiang, X., Islam, M., Morgan, J. & Doyle, M. P. Fate of *Listeria monocytogenes* in bovine manure – Amended soil. J. Food Prot. 67, 1676–1681 (2004).

30. Moura, A. et al. Whole genome-based population biology and epidemiological surveillance of *Listeria monocytogenes*. Nat. Microbiol. 2, 16185 (2016).

31. Kuenne, C. et al. Reassessment of the *Listeria monocytogenes* pan-genome reveals dynamic integration hotspots and mobile genetic elements as major components of the accessory genome. BMC Genomics 14, 47 (2013).

32. Cotter, P. D. et al. Listeriolysin S, a novel peptide haemolysin associated with a subset of lineage I Listeria monocytogenes. PLoS Pathog. 4, e1000144 (2008).

33. Lee, S., Ward, T. J., Jima, D. D., Parsons, C. & Kathariou, S. The arsenic resistanceassociated *Listeria* genomic island LGI2 exhibits sequence and integration site diversity and a propensity for three Listeria monocytogenes clones with enhanced virulence. Appl. Environ. Microbiol. 83, (2017).

34. Suchard, M. A. et al. Bayesian phylogenetic and phylodynamic data integration using BEAST 1.10. Virus Evol. 4, vey016 (2018).

35. Volz, E. M. & Frost, S. D. W. Scalable relaxed clock phylogenetic dating. Virus Evol. 3, 1–9 (2017).

36. Ishikawa, S. A., Zhukova, A., Iwasaki, W. & Gascuel, O. A fast likelihood method to reconstruct and visualize ancestral scenarios. Mol. Biol. Evol. 36, 2069–2085 (2019).

37. Bowling, G. A. The introduction of cattle into colonial North America. J. Dairy Sci. 25, 129–154 (1942).

38. Drummond, A. J., Rambaut, A., Shapiro, B. & Pybus, O. G. Bayesian coalescent inference of past population dynamics from molecular sequences. Mol. Biol. Evol. 22, 1185–1192 (2005).

39. Tajima, F. Statistical method for testing the neutral mutation hypothesis by DNA polymorphism. Genetics 123, 585–595 (1989).

40. World Trade Organization. World trade report. (World Trade Organization, 2013).

41. Zimmerman, W. D. Live cattle export trade between United States and Great Britain, 1868-1885. Agric. Hist. 36, 46–52 (1962).

42. Burroughs, W. J. & World Meteorological Organization. Climate: into the 21st century. (Cambridge University Press, 2003).

43. Groombridge, B. Global Biodiversity: status of the Earth’s living resources. (Springer Netherlands, 1992).

44. Franz, E. et al. Phylogeographic analysis reveals multiple international transmission events have driven the global emergence of *Escherichia coli* O157:H7. Clin. Infect. Dis. 69, 428–437 (2019).

45. Mourkas, E. et al. Agricultural intensification and the evolution of host specialism in the enteric pathogen *Campylobacter jejuni*. Proc. Natl. Acad. Sci. 117, 11018–11028 (2020).

46. LeBlanc, S. J., Lissemore, K. D., Kelton, D. F., Duffield, T. F. & Leslie, K. E. Major advances in disease prevention in dairy cattle. J. Dairy Sci. 89, 1267–1279 (2006).

47. Cartwright, E. J. et al. Listeriosis outbreaks and associated food vehicles, United States, 1998-2008. Emerg. Infect. Dis. 19, 1–9 (2013).

48. De Valk, H. et al. Two consecutive nationwide outbreaks of listeriosis in France, October 1999-February 2000. Am. J. Epidemiol. 154, 944–950 (2001).

49. Goulet, V. et al. Effect of prevention measures on incidence of human listeriosis, France, 1987-1997. Emerg. Infect. Dis. 7, 983–989 (2001).

50. Tappero, J. W., Schuchat, A., Deaver, K. A., Mascola, L. & Wenger, J. D. Reduction in the incidence of human listeriosis in the United States. Effectiveness of prevention efforts? JAMA 273, 1118–22 (1995).

51. Kathariou, S. Foodborne outbreaks of listeriosis and epidemic-associated lineages of *Listeria monocytogenes*. in Microbial Food Safety in Animal Agriculture 243–256 (Blackwell Publishing, 2008). doi:10.1002/9780470752616.ch25

52. Maury, M. M. et al. Spontaneous loss of virulence in natural populations of *Listeria monocytogenes*. Infect. Immun. 85, 1–13 (2017).

53. Castro, H., Jaakkonen, A., Hakkinen, M., Korkeala, H. & Lindström, M. Occurrence, persistence, and contamination routes of *Listeria monocytogenes* genotypes on three Finnish dairy cattle farms: A longitudinal study. Appl. Environ. Microbiol. 84, (2018).

54. Moura, A. et al. Real-time whole-genome sequencing for surveillance of *Listeria monocytogenes*, France. Emerg. Infect. Dis. 23, 1462–1470 (2017).

55. Criscuolo, A. & Brisse, S. AlienTrimmer: A tool to quickly and accurately trim off multiple short contaminant sequences from high-throughput sequencing reads. Genomics 102, 500–506 (2013).

56. Liu, Y., Schröder, J. & Schmidt, B. Musket: A multistage k-mer spectrum-based error corrector for Illumina sequence data. Bioinformatics 29, 308–315 (2013).

57. Andrews, S. FastQC: a quality control tool for high throughput sequence data. Available online at: http://www.bioinformatics.babraham.ac.uk/projects/fastqc (2010).

58. Bankevich, A. et al. SPAdes: a new genome assembly algorithm and its applications to single-cell sequencing. J. Comput. Biol. 19, 455–477 (2012).

59. Seemann, T. Prokka: Rapid prokaryotic genome annotation. Bioinformatics 30, 2068–2069 (2014).

60. Huerta-Cepas, J. et al. Fast genome-wide functional annotation through orthology assignment by eggNOG-mapper. Mol. Biol. Evol. 34, 2115–2122 (2017).

61. Buchfink, B., Xie, C. & Huson, D. H. Fast and sensitive protein alignment using DIAMOND. Nature Methods 12, 59–60 (2014).

62. Robertson, J. & Nash, J. H. E. MOB-suite: software tools for clustering, reconstruction and typing of plasmids from draft assemblies. Microb. genomics 4, (2018).

63. Arndt, D. et al. PHASTER: a better, faster version of the PHAST phage search tool. Nucleic Acids Res. 44, W16–W21 (2016).

64. Jolley, K. A. & Maiden, M. C. J. BIGSdb: Scalable analysis of bacterial genome variation at the population level. BMC Bioinformatics 11, 595 (2010).

65. Page, A. J. et al. Roary: Rapid large-scale prokaryote pan genome analysis. Bioinformatics 31, 3691–3693 (2015).

66. Collins, C. & Didelot, X. A phylogenetic method to perform genome-wide association studies in microbes that accounts for population structure and recombination. PLoS Comput. Biol. 14, e1005958 (2018).

67. Doumith, M., Buchrieser, C., Glaser, P., Jacquet, C. & Martin, P. Differentiation of the major *Listeria monocytogenes* serovars by multiplex PCR. J. Clin. Microbiol. 42, 3819–3822 (2004).

68. Dixon, P. VEGAN, a package of R functions for community ecology. J. Veg. Sci. 14, 927–930 (2003).

69. Edgar, R. C. MUSCLE: Multiple sequence alignment with high accuracy and high throughput. Nucleic Acids Res. 32, 113 (2004).

70. Mascola, L. et al. Listeriosis: an uncommon opportunistic infection in patients with acquired immunodeficiency syndrome. A report of five cases and a review of the literature. Am. J. Med. 84, 162–164 (1988).

71. Croucher, N. J. et al. Rapid phylogenetic analysis of large samples of recombinant bacterial whole genome sequences using Gubbins. Nucleic Acids Res. 43, e15 (2015).

72. Nguyen, L. T., Schmidt, H. A., Von Haeseler, A. & Minh, B. Q. IQ-TREE: A fast and effective stochastic algorithm for estimating maximum-likelihood phylogenies. Mol. Biol. Evol. 32, 268–274 (2015).

73. Tavaré, S. Some probabilistic and statistical problems in the analysis of DNA sequences. American Mathematical Society: Lectures on Mathematics in the Life Sciences (1986). doi:citeulike-article-id:4801403

74. Kalyaanamoorthy, S., Minh, B. Q., Wong, T. K. F., Von Haeseler, A. & Jermiin, L. S. ModelFinder: Fast model selection for accurate phylogenetic estimates. Nat. Methods 14, 587–589 (2017).

75. Minh, B. Q., Nguyen, M. A. T. & von Haeseler, A. Ultrafast approximation for phylogenetic bootstrap. Mol. Biol. Evol. 30, 1188–1195 (2013).

76. Yu, G., Smith, D. K., Zhu, H., Guan, Y. & Lam, T. T.-Y. ggtree: an r package for visualization and annotation of phylogenetic trees with their covariates and other associated data. Methods Ecol. Evol. 8, 28–36 (2017).

77. Letunic, I. & Bork, P. Interactive tree of life (iTOL) v3: an online tool for the display and annotation of phylogenetic and other trees. Nucleic Acids Res. 44, W242–-245 (2016).

78. Page, A. J. et al. SNP-sites: rapid efficient extraction of SNPs from multi-FASTA alignments. Microb. genomics 2, e000056 (2016).

79. Pfeifer, B., Wittelsbürger, U., Ramos-Onsins, S. E. & Lercher, M. J. PopGenome: an efficient Swiss army knife for population genomic analyses in R. Mol. Biol. Evol. 31, 1929–1936 (2014).

80. Rambaut, A., Lam, T. T., Max Carvalho, L. & Pybus, O. G. Exploring the temporal structure of heterochronous sequences using TempEst (formerly Path-O-Gen). Virus Evol. 2, (2016).

81. Drummond, A. J., Ho, S. Y. W., Phillips, M. J. & Rambaut, A. Relaxed phylogenetics and dating with confidence. PLoS Biol. 4, e88 (2006).

82. Rambaut, A., Drummond, A. J., Xie, D., Baele, G. & Suchard, M. A. Posterior summarization in Bayesian phylogenetics using Tracer 1.7. Syst. Biol. 67, 901–904 (2018).

83. Firth, C. et al. Using time-structured data to estimate evolutionary rates of double-stranded DNA viruses. Mol. Biol. Evol. 27, 2038–51 (2010).

84. Didelot, X., Croucher, N. J., Bentley, S. D., Harris, S. R. & Wilson, D. J. Bayesian inference of ancestral dates on bacterial phylogenetic trees. Nucleic Acids Res. 46, e134 (2018).

85. Himmelmann, L. & Metzler, D. TreeTime: an extensible C++ software package for Bayesian phylogeny reconstruction with time-calibration. Bioinformatics 25, 2440–1 (2009).

86. Paradis, E. & Schliep, K. ape 5.0: an environment for modern phylogenetics and evolutionary analyses in R. Bioinformatics 35, 526–528 (2019).

87. Salje, H. et al. Dengue diversity across spatial and temporal scales: Local structure and the effect of host population size. Science (80-.). 355, 1302–1306 (2017).

